# A comprehensive temporal patterning gene network in *Drosophila* medulla neuroblasts revealed by single-cell RNA sequencing

**DOI:** 10.1101/2021.06.12.448145

**Authors:** Hailun Zhu, Sihai Dave Zhao, Alokananda Ray, Yu Zhang, Xin Li

## Abstract

During development, neural stem cells are temporally patterned to sequentially generate a variety of neural types before exiting the cell cycle. Temporal patterning is well-studied in *Drosophila*, where neural stem cells called neuroblasts sequentially express cascades of Temporal Transcription Factors (TTFs) to control the birth-order dependent neural specification. However, currently known TTFs were mostly identified through candidate approaches and may not be complete. In addition, many fundamental questions remain concerning the TTF cascade initiation, progression, and termination. It is also not known why temporal progression only happens in neuroblasts but not in their differentiated progeny. In this work, we performed single-cell RNA sequencing of *Drosophila* medulla neuroblasts of all ages to study the temporal patterning process with single-cell resolution. Our scRNA-seq data revealed that sets of genes involved in different biological processes show high to low or low to high gradients as neuroblasts age. We also identified a list of novel TTFs, and experimentally characterized their roles in the temporal progression and neural fate specification. Our study revealed a comprehensive temporal gene network that patterns medulla neuroblasts from start to end. Furthermore, we found that the progression and termination of this temporal cascade also require transcription factors differentially expressed along the differentiation axis (neuroblasts -> -> neurons). Lola proteins function as a speed modulator of temporal progression in neuroblasts; while Nerfin-1, a factor required to suppress de-differentiation in post-mitotic neurons, acts at the final temporal stage together with the last TTF of the cascade, to promote the switch to gliogenesis and the cell cycle exit. Our comprehensive study of the medulla neuroblast temporal cascade illustrated mechanisms that might be conserved in the temporal patterning of neural stem cells.

## Introduction

The heterogeneity of neural fates builds the foundation for constructing complex neural circuits. Integration of spatial and temporal patterning of neural stem cells allows neural progeny to adopt a spectrum of identities [reviewed in ^1–4^]. Spatial patterning specifies distinct lineages, while within a specific lineage, temporal patterning further expands the neural diversity, as neural stem cells undergo gradual transitions along with self-renewal and give rise to a successive series of neural fates. A molecular mechanism of temporal patterning was identified in the nervous system of *Drosophila*. In *Drosophila*, neural stem cells called neuroblasts (NBs), were found to sequentially express certain cascades of temporal transcription factors (TTFs) and other temporal regulators that specify neural fates [reviewed in ^5, 6^]. The first TTF cascade was identified in the embryonic ventral nerve cord, where Hunchback (Hb), Kruppel (Kr), Nubbin/Pdm2 (Pdm), Castor (Cas) and Grainy head (Grh) are sequentially expressed in NBs as they age, and are required for the sequential specification of different neural fates^7–12^. Postembryonic NBs, including larval ventral nerve cord, central brain and optic lobe NBs, also employ temporal cascades that are yet different from the embryonic one ^13–20^. In addition, the intermediate neural progenitors (INPs) of type II NBs are also temporally patterned by a TTF cascade to further expand the neural diversity ^21–23^. In vertebrates, there is also accumulating evidence that neural stem cells undergo TTF dependent temporal patterning [reviewed in ^24, 25^]. For example, a number of transcription factors were shown to function in retinal progenitors or cortical progenitors to regulate temporal specification of neural fates ^26–35^. Recently, single-cell transcriptomics studies of retinal progenitors and cortical progenitors revealed age-dependent dynamic changes in transcriptional profiles that are transmitted to their progeny ^28, 36^. These studies together suggest that TTF-dependent temporal patterning might be a general mechanism.

The model system we use to study temporal patterning is the medulla part of the *Drosophila* optic lobe. During development, a wave of neurogenesis sweeps from medial to lateral in the outer proliferation center (OPC) and sequentially converts symmetrically-dividing neuroepithelial cells (NE) into medulla NBs ^37–39^. Medulla NBs divide asymmetrically multiple times to make a series of Ganglion Mother Cells (GMCs) which then divide to produce postmitotic progeny. Due to the spreading of the neurogenesis wave, neuroblasts of different ages and their progeny are orderly aligned on the medial to lateral spatial axis in the developing larval brain (Figure 1A). This feature makes medulla a great system to study temporal patterning. Previous studies showed that medulla NBs sequentially express Homothorax (Hth), Kumpfuss (Klu), Eyeless (Ey), Sloppy paired (Slp), Dichaete (D) and Tailless (Tll) as they age ^18, 19^. Among them, Hth, Ey, Slp and D are each required for the expression of the corresponding neuronal transcription factors to control neural fates, but loss of Klu caused a NB proliferation defect, precluding examination of neural fates ^18, 19, 40^. Similar to vertebrate retinal and cortical progenitors, medulla NBs switch to gliogenesis at the end of the lineage and then exit the cell cycle ^18, 41^.

**Figure 1.**
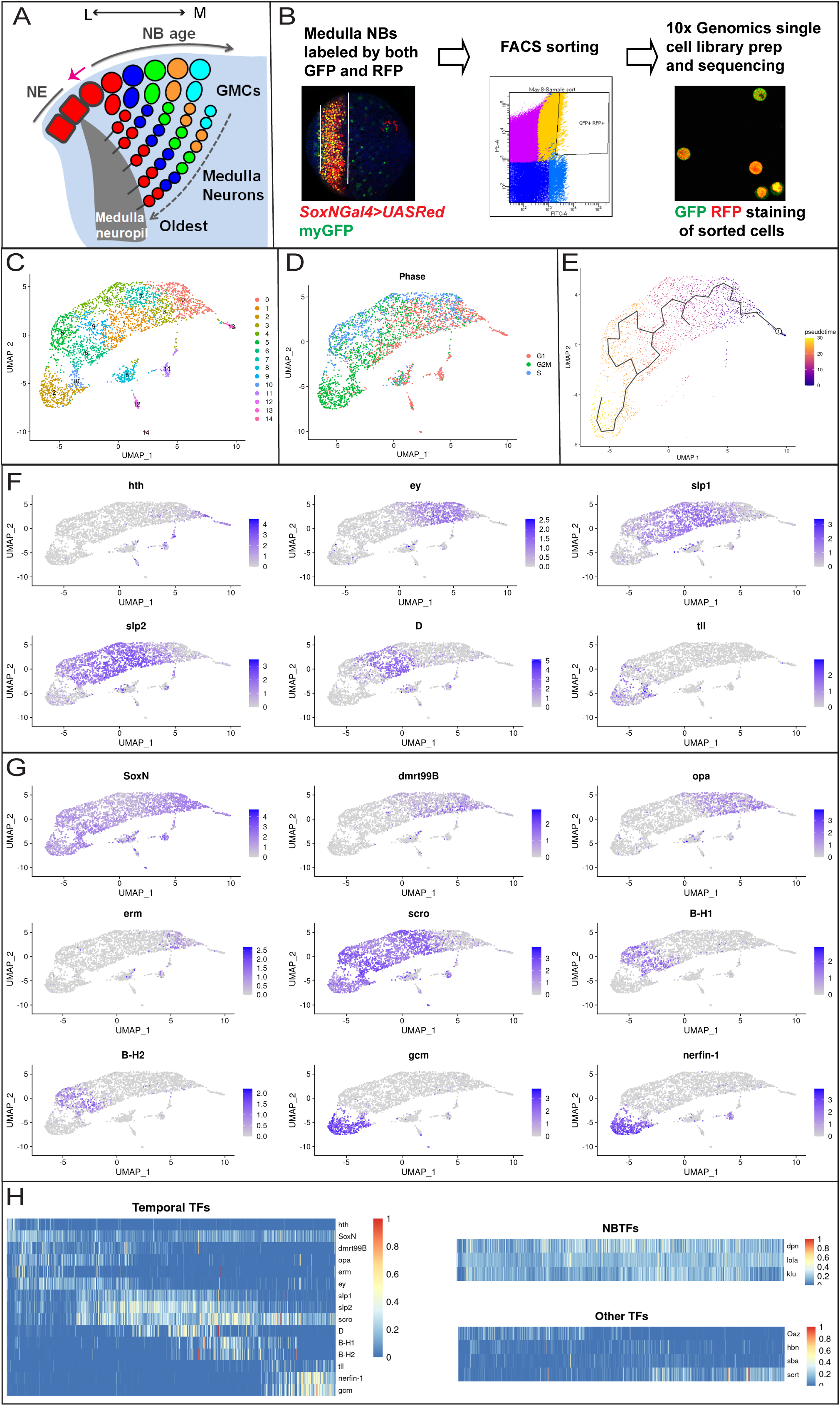
ScRNA-seq of *Drosophila* medulla neuroblasts. (A) A schematic drawing of the developing *Drosophila* medulla at the 3^rd^ instar larval stage. A neurogenesis wave (pink arrow) spreads from medial (M) to lateral (L), and sequentially converts NE cells into NBs. Thus, NBs from the youngest to the oldest are aligned on the lateral to medial axis. NBs sequentially express different TTFs, and generate differently fated progeny. The earliest-born neurons of each NB lineage are located closest to the medulla neuropil. The later-born neurons are located at more and more superficial layers. (B) The strategy and work-flow of the scRNA-seq of FACS sorted medulla NBs. Medulla NBs are uniquely labeled by the combination of *SoxNGal4>UAS-RedStinger* (red) and *E(spl)mγGFP* (green). Dissected larval brains were dissociated into single-cell suspension and subjected to FACS sorting to enrich medulla NBs. Then 10x V3 Single Cell libraries were generated using the sorted cells, and sequenced. (C) The sequenced cells were partitioned into 15 clusters using Seurat and visualized on UMAP plots. (D) The estimated cell cycle phase of each sequenced cell was visualized on UMAP plots. (E) Pseudotime trajectories generated using Monocle3, with the purple color stands for the earliest pseudotime, and yellow color stands for the latest pseudotime. (F) The expression patterns of known TTFs (Hth, Ey, Slp1, Slp2, D and Tll) verify the pseudotime trajectories. (G) The expression patterns of newly identified TTFs visualized on UMAP plots. (H) Heatmaps showing the expression levels of three classes of TFs across the pseudotime. Temporal TFs include known and novel TTFs; NBTFs include three TFs expressed in NBs of all ages; Other TFs include four TFs that show a temporal expression profile, but were not further characterized. The expression levels are visualized as a percentage of the maximum observed expression across all cells.

In the medulla TTF cascade, there are still fundamental questions remaining. First, the several TTFs identified through candidate antibody screening may not compose the complete TTF sequence. Second, while Ey, Slp and D are each required for the next TTF to be activated, no cross-regulation was identified among Hth, Klu and Ey. Thus, it is not known how the Ey->Slp->D->Tll TTF cascade is initiated. Third, it was not clear how the oldest medulla NBs switch to gliogenesis, end the temporal progression and exit the cell cycle. Finally, an even broader question concerns the regulation of the temporal cascade. As a previous TTF is only necessary but not sufficient to activate the next TTF in medulla NBs, additional regulators and molecular mechanisms might be involved to achieve the proper regulation of the cascade. Furthermore, temporal cascade progression only happens in neuroblasts but not in their differentiated progeny, which makes us to question whether genes functioning in the differentiation axis (NBs -> GMCs -> neurons) might play a role in regulating the temporal progression, which has never been studied before.

To discover all unknown TTFs and additional regulators, as well as to get a global view of the dynamic temporal patterning process of medulla neuroblasts, we applied single cell RNA sequencing (scRNA-seq) to our model system. ScRNA-seq is increasingly used as an unbiased approach to characterize heterogeneous tissues, including invertebrate and vertebrate nervous systems [reviewed in ^42–45^]. The *Drosophila* medulla represents a great system to study temporal patterning using scRNA-seq, because at a single time point during development, we can obtain a continuous population of neuroblasts of all ages, and furthermore, we can use known TTFs to mark the neuroblast age and verify the inferred pseudotime trajectory. Applying scRNA-seq to the *Drosophila* medulla neuroblasts enabled us to capture all temporal stages and to reveal the gradual changing of neuroblast transcriptome with single-cell-cycle resolution. We observed growth and metabolism related genes showed high to low gradients, while genes involved in gene expression regulation and neural differentiation showed low to high gradients of expression as neuroblasts age. We identified a list of novel TTF candidates, among which, SoxNeuro (SoxN), doublesex-Mab related 99B (Dmrt99B), Odd paired (Opa), Earmuff (Erm), Scarecrow (Scro), BarH1, BarH2 and Glial cells missing (Gcm) were confirmed to be required for the normal temporal patterning of medulla neuroblasts. We identified extensive cross-regulations among these novel TTFs and known TTFs, that generally follow the rule that a TTF is required to activate the next TTF and repress the previous TTF, but with important exceptions and complexities. Our study revealed a comprehensive temporal patterning cascade: Hth + SoxN + dmrt99B -> Opa -> Ey+Erm -> Ey+Opa -> Slp+Scro -> D -> B-H1&2->Tll, Gcm, that controls the sequential generation of different neural types by regulating the expression of specific neuronal transcription factors. Finally, Gcm but not Tll is required for both the transition from neurogenesis to gliogenesis and the cell cycle exit. Moreover, in pursuit of the mechanism behind the regulation of the temporal cascade, we found that the progression and termination of the TTF temporal cascade require genes differentially expressed along the differentiation axis (NBs -> GMCs -> neurons) including *lola* and *nerfin-1*, providing clues as to why the temporal progression only proceeds in neural stem cells.

## Results

### ScRNA-seq revealed the dynamic transcriptome of *Drosophila* medulla neuroblasts as they age

To perform single-cell transcriptional profiling of *Drosophila* medulla neuroblasts, we dissociated dissected larval brains into single cell suspension and used fluorescence-activated cell sorting (FACS) to enrich medulla neuroblasts (Figure 1B). Medulla neuroblasts were sorted by the co-expression of two transgenes, *SoxN*Gal4 driving UASRed that is expressed in all medulla cells, and E(spl)mγ::GFP that is expressed in all neuroblasts. We performed two rounds of scRNA-seq of sorted medulla neuroblasts using the 10x Genomics Chromium platform. We combined data from both sequencing experiments into a single analysis by integrating them using Seurat ^46^. After quality control and filtering for neuroblasts, our data contained 3074 cells expressing between 261 and 6409 genes, with a median of 3682 expressed genes per cell.

To characterize the developmental states of sequenced neuroblasts, we used Seurat to partition the cells into 15 clusters, which we visualized on two-dimensional uniform manifold approximation and projection (UMAP)^47^ plots (Figure 1C). Among them, clusters 8, 11, 12, and 14 appear to be outliers, while all other clusters form a continuous cell stream, representing the main body of medulla neuroblasts. We also used Seurat to estimate the cell cycle phase of each cell (Figure 1D), based on the expression of known *Drosophila* cell cycle genes from Tinyatlas at Github (Supplementary Table 1). In the main body of medulla neuroblasts, cluster 13 contains mostly G1 cells, while cluster 2 contains mostly G2/M cells, and in other clusters cells of different phases are mixed. Next we inferred single-cell pseudotime trajectories using Monocle3 ^48–50^ (Figure 1E). For this analysis, we removed the outlying clusters 8, 11, 12, and 14. The inferred developmental pseudotime appears to progress from right to left. Finally, the expression patterns of known TTFs in the medulla NB temporal cascade, in the order of Hth, Ey, Slp1/Slp2, D and Tll, validated the inferred pseudotime trajectory of neuroblasts (Figure 1F). These data confirmed that our scRNAseq captured a continuous population of neuroblasts of all ages that formed a chronotopic map, with cluster 13 representing the newly-born NBs at G1 phase, cluster 0 containing the youngest neuroblasts that have started proliferation, and cluster 2 containing the oldest neuroblasts undergoing the terminal division (mostly at G2/M phases).

### Two sets of genes show opposite expression gradients as medulla NBs age

Among the genes whose expression change with pseudotime, there are two sets of genes with high to low or low to high gradients, respectively. Genes encoding ribosomal proteins and metabolism enzymes showed significant high to low gradients, while genes involved in gene expression regulation and neural development showed low to high gradients. Negative regulators of cell growth and proliferation are gradually upregulated at later stages (Figure S1A,B) These data suggest that as medulla NBs age, the cell growth is gradually downregulated, while differentiation related genes are up-regulated.

### DEG analysis identified Candidate novel TTFs

With the neuroblast transcriptome profile of all ages, we sought to identify more potential TTFs that had temporal expression patterns. Known TTFs were used to mark the relative age of a neuroblast, and indicate the position where a potential TTF might stand in the temporal cascade. We obtained a pre-compiled list of 755 *Drosophila* TFs^51^ identified based on bioinformatic analysis and manual curation. For each non-outlying cell cluster, we performed the DEG (Differentially Expressed Genes) analysis and identified the top 10 TFs in this list that were differentially expressed between the clusters (Figure S1C). We next examined the expression of these TFs at the protein level by immunostaining of 3^rd^ instar larval brains using available antibodies or GFP-fusion lines. To determine whether these TFs play crucial roles in the medulla temporal cascade, we examined cross-regulations between the novel TFs and known TTFs. We also assessed whether these TFs are involved in progeny fate specification. After screening through these TFs, we identified SoxN, Dmrt99B, Opa, Erm, Scro, BarH1, BarH2 and Gcm as novel TTFs in the temporal cascade (Figure 1G, H). According to the scRNA-seq data, SoxN transcripts are already present in the newly-born NBs similar to Hth, remain high in the youngest NBs, and then become reduced from the Ey stage to the D stage and increase again after the D stage; Dmrt99B transcripts are present from the newly-born NBs until the Slp stage NBs; the expression of Opa transcripts starts in the youngest NBs before Ey, and continues until the end of the Ey stage, but with a small gap in the middle corresponding to the early Ey stage; Erm transcripts are present at this gap between the two groups of opa-expressing NBs, i.e. in the early Ey stage NBs; Scro transcripts are present in NBs from about the Slp stage to the final stage; BarH1 and BarH2 transcripts are present in a group of NBs older than the D stage NBs but younger than the Tll stage NBs; Gcm transcripts are present in the NBs of the final stage, later than the Tll stage (Figure 1G,H). For a few other candidate TFs with temporal expression patterns, including Oaz, Hbn, Scrt and Sba (Figure 1H), we either did not observe an effect on the temporal progression (Oaz, Sba, Scrt) using available RNAi lines, or we are lacking effective reagents (Hbn). Therefore, we did not include these TFs in further analysis.

### SoxN and Dmrt99B cooperate with Hth to specify the first temporal identity

According to our scRNA-seq data, SoxN transcripts have a rather broad distribution in NBs. However, antibody staining showed that SoxN protein is only expressed before the Ey stage (Figure 2A-A’’), indicating that special post-transcriptional regulation of SoxN exists at later stages. SoxN is a SoxB family HMG-domain transcription factor involved in neuroblast formation and neuron differentiation in the ventral nerve cord^52–55^. Its expression has been noted in the medulla^19^, but its function in the medulla has not been studied.

**Figure 2.**
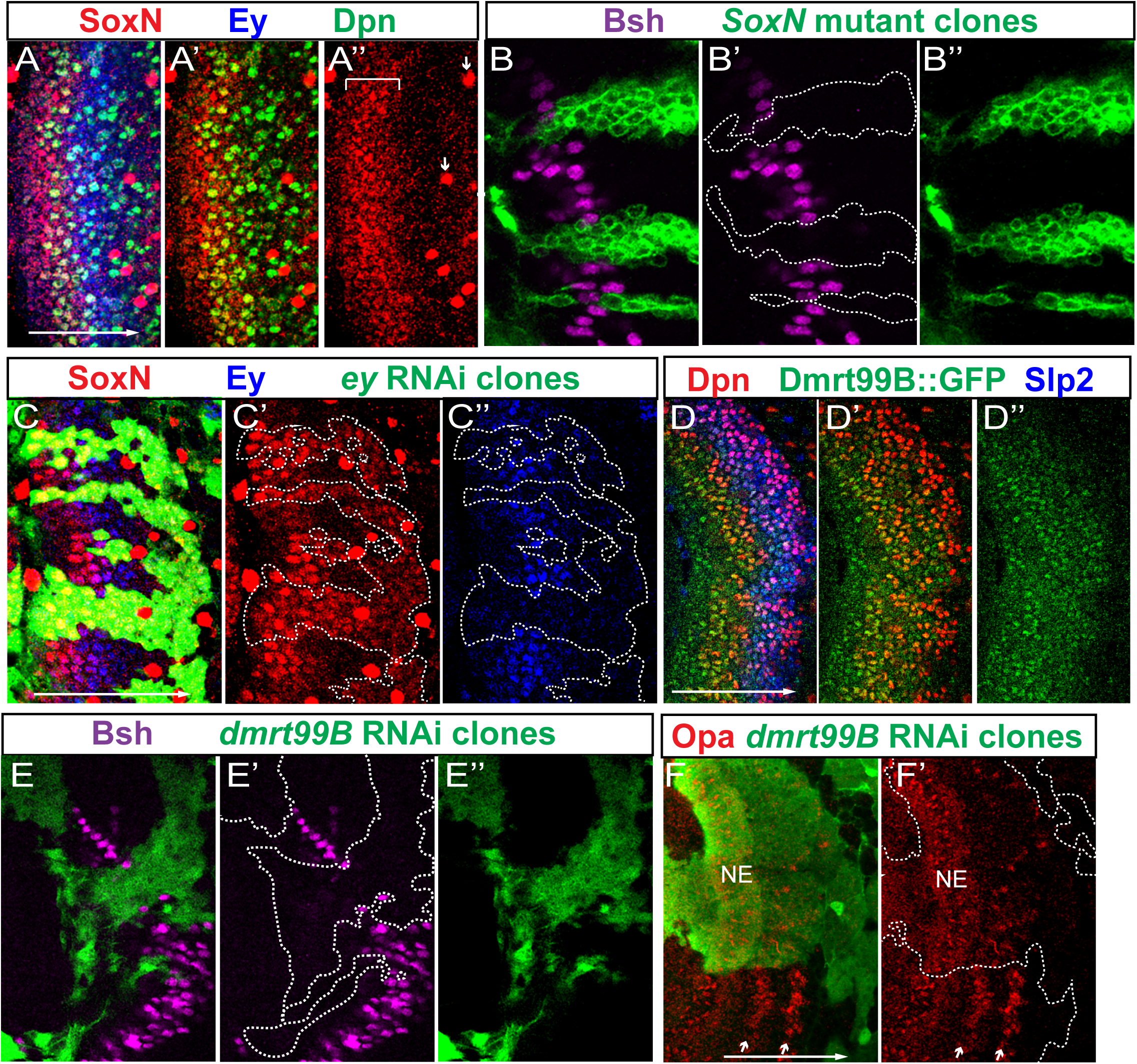
SoxN and Dmrt99B are two TTFs that function together with Hth in the first temporal stage. In all images of this and the following figures, lateral is to the left, and medial is to the right. The large white arrow from left to right indicates the NB age from the youngest to oldest. White dashed lines indicate clone margin unless otherwise noted. (A-A’’) SoxN protein (red) is expressed before NB formation and in the youngest NBs (marked by Dpn in green) before Ey (blue), and this expression domain is indicated by a white bracket. SoxN is also expressed in glial cells, and examples are indicated by small white arrows. (B-B’’) In *SoxN^NC14^* mutant clones marked by GFP (green), Bsh (magenta) is lost (in 12 out of 12 clones). (C-C’’) In *eyRNAi* clones marked by GFP (green), Ey (blue) is lost, and SoxN (red) expands into older NBs (in 11 out of 11 clones). (D-D’’) Dmrt99B::GFP (green) is expressed before NB formation and in the young NBs (NBs marked by Dpn in red) until early Slp (blue) stage. (E-E’’) In *Dmrt99B-RNAi* clones marked by GFP (green), Bsh (magenta) is lost (in 7 out of 7 clones). (F,F’) In *Dmrt99B-RNAi* clones marked by GFP (green), Opa (red) is lost or delayed (4 out of 4 clones). White arrows indicate the two stripes of Opa expression in NBs.

Similar to Hth, SoxN expression starts in NE cells before they are transformed into NBs marked by Deadpan (Dpn), and continues in the youngest NBs until the Ey stage (Figure 2A-A’’). SoxN is also inherited in GMCs and neurons generated before the Ey stage (Figure S2A,A’). To test if SoxN is required for neuron fate specification, we generated *SoxN* homozygous mutant clones in otherwise heterozygous brains using the MARCM system^56^. First we observed that SoxN staining is lost in *SoxN* mutant clones, validating the specificity of the antibody^55^ and the mutant (Figure S2B,B’). In the mutant clones, Bsh (brain-specific homeobox) expressing neurons which are generated in the Hth stage ^18, 19, 57^ were lost, suggesting that SoxN cooperates with Hth to specify the Bsh neuron fate (Figure 2B-B’’). Moreover, with loss of SoxN, Hth expression in NBs was not affected, and the following temporal cascade still proceeded as indicated by the Ey and Slp expression in mutant NBs (Figure S2C-D’’’). These results suggest that SoxN is not required for the initiation of the temporal gene cascade in NBs. To test if the termination of SoxN is due to the cascade progression, we generated *ey* RNAi clones using the *ay*Gal4 system^58^ as well as *slp* mutant MARCM clones. SoxN expression was expanded into older NBs in *ey* RNAi clones but not affected in *slp* mutant clones, suggesting that Ey inhibits the expression of SoxN (Figure 2C-C’’, Figure S2E-E’’). Therefore, SoxN is another TTF that determines the first temporal stage together with Hth. However, neither SoxN nor Hth is required for the progression of the cascade.

Another TF gene expressed in the youngest NBs is *dmrt99B*, encoding a DMRT (doublesex- and mab-3-related) transcription factor ^59, 60^. A Dmrt99B::GFP fusion protein expressed under genomic BAC endogenous control (from modERN Project) is turned on before the NE to NB transition and remains high in young NBs, and decreases at the beginning of the Slp stage (Figure 2D-D’’). In *dmrt99B* RNAi clones generated using the *ay*Gal4 system, the NE to NB transition is not affected, but the Bsh expression is lost (Figure 2E-E’’, Figure S2G,G’), suggesting that Dmrt99B is also required for the first temporal fate. In contrast to SoxN and Hth, Dmrt99B is required for the later TTF cascade. In *dmrt99B* RNAi clones, the expression of the next TTF Opa (see the section below) is lost or dramatically delayed (Figure 2F,F’), Slp2 is also delayed and D is lost (Figure S2H-H’’’).

In summary, SoxN, Dmrt99B and Hth are three TTFs that are first turned on in the NE and all of them are required for the first temporal fate, but only loss of Dmrt99B affected the proper progression of the subsequent temporal cascade (Figure 3L). However, it is possible that partial redundancy may exist within these three TTFs.

**Figure 3.**
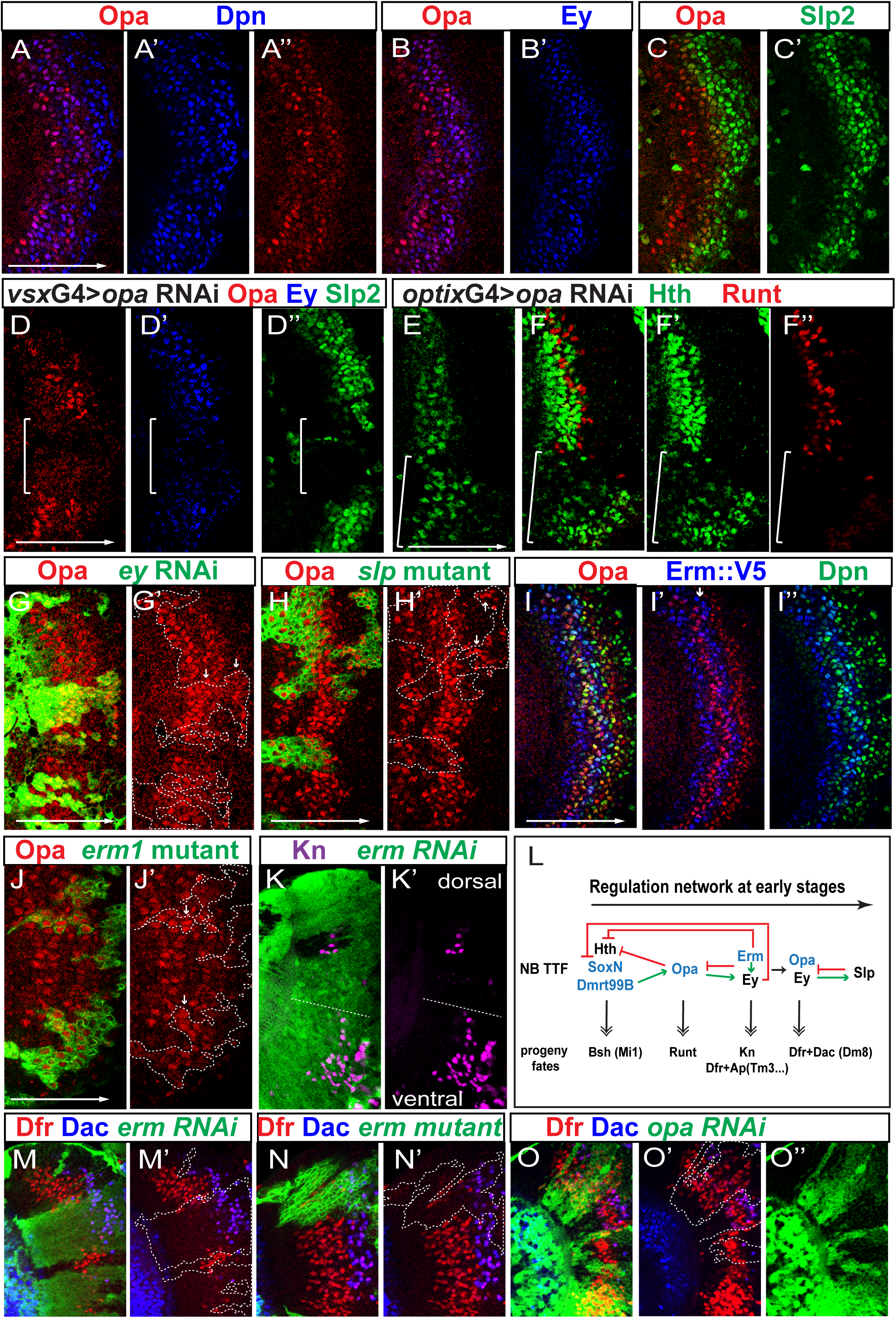
Opa and Erm are among the early TTFs. (A-A’’) Opa protein (red) is expressed in two stripes of NBs (marked by Dpn in blue). (B-B’) The first stripe of Opa (red) is downregulated as Ey (blue) is upregulated, the second stripe of Opa overlaps with the Ey stripe. (C-C’) The second stripe of Opa is downregulated as Slp (green) is upregulated. (D-D’’) When *opaRNAi* is driven by *vsxGal4*, Opa (red) is lost in the center domain (cOPC, white bracket), Ey (blue) and Slp2 (green) are also mostly lost in the same domain (in 11 out of 11 brains). (E-F’’) *opaRNAi* is driven by *optixGal4* in mOPC indicated by a white bracket. (E) At a surface focal plane, Hth (green) is expanded into older NBs in mOPC (in 3 out of 3 brains). (F-F’’) At a deeper progeny focal plane, Hth expression (green) is expanded into later-born progeny, and Runt (red) expressing neurons are lost (in 3 out of 3 brains). (G,G’) In *ey-RNAi* clones marked by GFP (green), Opa expression (red) is de-repressed in the gap and also expanded into older NBs (in 15 out of 15 clones). (H,H’) In *slp* mutant clones marked by GFP (green), Opa expression (red) is expanded into older NBs (3 out of 3 clones). (I-I’’) Erm::V5 protein (blue) is expressed in two stripes, one before the NB (marked by Dpn in green) formation, and the other (white arrow) is between the two Opa stripes (red). (J,J’) In *erm^1^* mutant clones marked by GFP (green), Opa expression (red) is de-repressed in the gap (white arrows) (in 13 out of 13 clones). (K,K’) In *erm-RNAi* clones marked by GFP (green), Kn (magenta) expressing neurons are lost on the dorsal side (in 15 out of 15 clones), but still present on the ventral side (in 13 out of 14 clones). A white dashed line separates the dorsal vs ventral side. (L) A schematic showing the regulatory network among early TTFs and the neuron fates generated at each stage. (M-M’) In *erm-RNAi* clones marked by GFP (green), neurons expressing Dfr (red) but not Dac (blue) are lost, and the remaining neurons express both Dfr and Dac thus appear purple (in 13 out of 15 clones). (N,N’) In *erm^1^* mutant clones marked by GFP (green), neurons expressing Dfr (red) but not Dac (blue) are lost, and the remaining neurons express both Dfr and Dac thus appear purple (in 11 out of 13 clones). (O-O’’) In *opa-RNAi* clones marked by GFP (green), neurons expressing both Dfr (red) and Dac (blue) are lost, but neurons expressing only Dfr are expanded (in 8 out of 8 clones).

### Opa is the missing link between the NE TTFs and the NB TTF cascade

Opa is a zinc finger transcription factor homologous to mammalian Zic proteins, and has been shown to function in the embryonic patterning process as a pioneer factor ^61, 62^, as well as in adult head development^63, 64^. In the type II neuroblast lineage, Opa is expressed in early-born INPs and required for the temporal progression of the INP temporal cascade^23^.

We examined Opa protein expression in the medulla, and observed that it is expressed in two stripes of NBs, consistent with the transcript pattern (Figure 3A-A’’). Opa is not expressed in NE cells, and its expression is activated at the same time as Dpn in the youngest NBs (Figure 3A-A’’). The first Opa stripe is more towards the lateral compared to the Ey stripe (i.e. the first wave of Opa expression in NBs is before Ey), while the second stripe is overlapping with the medial half of the Ey stripe (i.e. the second wave of Opa expression is in late Ey stage NBs) (Figure 3B,B’). The second stripe of Opa expression terminates as Slp expression reaches its peak (Figure 3C,C’). Inside the medulla, two layers of Opa-expressing neurons, seemingly born from the two Opa windows, are observed. The neurons eventually stop expressing Opa during maturation (Figure S3A-A’’).

To test whether Opa is required in the temporal cascade, we generated *opa* mutant clones. With loss of Opa, Ey expression and the subsequent temporal cascade, as indicated by Slp2 expression, are greatly delayed (Figure S3B-B’’). The reason for a delay but not a complete loss of Ey expression might be that the clones are not generated early enough to completely eliminate Opa expression. To circumvent this problem, we used *vsx*G4, which is expressed in the central compartment of the medulla crescent (cOPC)^65^ (Figure S3L), to drive *opa* RNAi eliminating Opa expression from the start. In the region where Opa expression was absent, Ey expression and Slp2 expression were mostly lost (Figure 3D-D’’), suggesting that Opa is necessary for the temporal cascade to normally proceed towards the Ey stage. We also used another regional Gal4, *optix*G4, which is expressed in the main arms of the medulla crescent (mOPC regions) (Erclik et al., 2017) (Figure S3L) to drive opa RNAi and obtained the same phenotype. Meanwhile, in the region with *opa* RNAi, Hth expression was expanded in both NBs and progeny (Figure 3E, F,F’). Therefore, Opa is required to repress the previous TTF Hth, and activate the next TTF Ey.

Next, we examined whether known TTFs regulate *opa* expression. In *ey* RNAi clones, Opa expression persisted without a gap from the youngest NBs to the oldest NBs (Figure 3G,G’), suggesting that Ey (and/or any later TTF in the temporal cascade) is required to repress Opa in both the gap and the older NBs. In addition, neurons born at different time points all inherit Opa in the *ey* RNAi clones. Since the termination of the second stripe of Opa corelates with the upregulation of Slp, and Ey is required to activate Slp, it is possible that the de-repression of Opa in older NBs in *ey* RNAi clones is due to loss of Slp. To test whether Slp is required to terminate the second wave of Opa expression, we generated *slp* mutant clones, inside which the expression of the second stripe of Opa expanded towards the oldest NBs (Figure 3H,H’). However, the gap between the two stripes was not affected in *slp* mutant clones.

Finally, we examined whether Opa controls the specification of neural fates. A layer of neurons expressing Runt are born between the Hth stage and the Ey stage^19^ (Figure S3C). Runt expressing neurons are unaffected in *hth* mutant clones, and they are expanded in *ey* mutant clones, suggesting that a TTF repressed by Ey is responsible for the generation of Runt neurons (Figure S3D, E, E’). Consistently, over-expression of Ey in NBs is sufficient to block the production of Runt neurons (data not shown). Klu was implicated as a TTF specifying Runt neurons because over-expression of Klu led to extra Runt neurons^19^. However, Klu and Ey do not regulate the expression of each other, and *klu* mutant caused ectopic NB phenotype, precluding examination of neural fates^19^. Since the first stripe of Opa is turned on after Hth and before Ey, and we have shown that Ey is required to repress Opa, we examined whether Opa is required to specify Runt neuron fate. In *opa* RNAi regions, Runt neurons are largely lost (Figure 3F,F’’), suggesting that the first stripe of Opa indeed serves as a TTF between the Hth stage and the Ey stage, and is required for Runt neuron determination (Figure 3L).

### Erm is required to generate the two-stripe expression pattern of Opa

According to the scRNA-seq data, Erm transcripts are present at the gap between the two groups of Opa-expressing NBs. Erm, an ortholog of mammalian Fezf2, is a zinc-finger transcription factor shown to be expressed in INPs of type II neuroblast lineages where it maintains the INPs’ restricted developmental potential^66–68^. A role for Erm in temporal patterning has not been studied before. We used an Erm::V5 line that recapitulates the true expression of Erm^69^ to examine the protein expression pattern of Erm in the medulla. The staining for V5 marker showed that Erm is indeed expressed at the gap between the two Opa-expressing stripes in NBs, consistent with the mRNA pattern (Figure 3I-I’’). In addition, Erm is also inherited in the progeny located between the two layers of Opa^+^ neurons (Figure S3F). Interestingly, there is another stripe of Erm expressed in NE cells adjacent to the youngest NBs (Figure 3I-I’’). The expression pattern of Erm in NBs suggests that it might repress Opa expression and generate the gap. To address this hypothesis, we generated *erm^1^* mutant clones, and showed that with loss of Erm, the gap in Opa expression in NBs and progeny was no longer present (Figure 3J,J’, Figure S3G,G’), suggesting that Erm does function to repress Opa at the gap, possibly through cooperation with Ey. In *erm* mutant clones, Ey expression became weaker but was still present, and Hth expression was expanded (Figure S3H-I’). In *erm* RNAi clones, Opa showed the same de-repression phenotype (data not shown). In summary, we showed that Erm is activated in NBs at a similar time as Ey, and is required to repress Opa to generate the gap in Opa expression.

### The Ey stage is divided into two sub-temporal stages by Erm and the second stripe of Opa

The expression patterns and cross-regulations between Erm, Opa and Ey suggest that the Ey stage can be subdivided into two sub-temporal stages, with early Ey stage NBs co-expressing Ey and Erm, and late Ey stage NBs co-expressing Ey and the second stripe of Opa (Figure 1F,G,H, Figure 3B,B’,I,I’). It is possible that Ey stage NBs generate different neural types in these two sub-temporal stages. To test this hypothesis, we set out to examine whether loss of *erm* or *opa* affects the Ey progeny fates. As previously reported^18^, each medulla GMC divides to generate a Notch-on neuron which expresses Apterous (Ap) and a Notch-off neuron which doesn’t express Ap. The Notch-off neurons generated in the Ey stage inherit Ey expression, and they also express a bHLH transcription factor Knot (Kn) (Figure S3J-J’’). Ey is required for Kn expression, because Kn is lost in *ey* RNAi clones (Figure S3K,K’). Drifter (Dfr, also known as Vvl) is another transcription factor expressed in the Ey stage progeny, and is lost in *ey* mutant or RNAi clones^18, 19^. Dfr expressing neurons can be divided into two large populations. The first population is a layer of Notch-on neurons generated in the early Ey stage that express both Dfr and Ap (Figure S3 M-N’, Dfr^+^ cells between the two white dashed lines), and among this population some neurons also express a weak level of Dac, thus expressing all three TFs (white arrows in Figure S3 M-N’). The second population includes clusters of later-born Notch-off neurons that express both Dfr and Dac, but not Ap^18, 19, 57^ (Figure S3M’,N’, purple cells enclosed by green dashed lines). The Dfr^+^ Notch-on neurons are specified into several neural types including Tm3, Tm9, Mi10 etc^57^. The Dfr^+^ Dac^+^ Notch-off neurons are only generated in certain spatial domains, and they are specified as Dm8 multi-columnar neurons^57, 70^. These Dfr^+^ Dac^+^ N-off neurons are still produced in *slp* mutant clones, suggesting that they are born before the Slp stage (Figure S3O,O’).

Using these markers, we examined whether Erm is required for the specification of neural fates. In *erm* RNAi clones, Kn expressing neurons are always lost on the dorsal side of the medulla, but still present on the ventral side (Figure 3K,K’); the first population of Dfr expressing neurons are largely lost, but the second population neurons expressing both Dfr and Dac generated at a later stage are still present (Figure 3M,M’). In *erm* mutant clones, we observed the same phenotype (Figure 3N,N’). These data suggest that Erm is required for the generation of the first population of Dfr^+^ neurons in the early Ey stage but not for the second population (Figure 3L). Erm is also required for the production of Kn expressing neurons in the dorsal medulla, but Kn expressing neurons are still present in the ventral medulla with loss of Erm, and a possible reason is that another population of Kn^+^ neurons not dependent on Erm are generated in the ventral medulla only, possibly in the late Ey stage.

Next we examined the expression of Kn, Dfr and Dac in *opa* RNAi clones. Since Opa is required for Ey expression, and Ey is required for Kn and Dfr expression, we would expect that Kn and Dfr are lost in *opa* RNAi clones. However, only Kn is lost in *opa* RNAi clones (Figure S3P,P’), while Dfr expression is not lost and even expanded, suggesting that Dfr expression does not require Ey if Opa is not present. However, neurons expressing both Dfr and Dac are lost in *opa* RNAi clones (Figure 3O-O’’). These together suggest that Opa normally represses the generation of the first population Dfr^+^ neurons (Dfr^+^ Ap^+^ neurons), but is required for the generation of Dfr^+^ Dac^+^ neurons (second population). Erm and Ey together are required to turn off Opa at the early Ey stage to allow for the generation of Dfr^+^ Ap^+^ neurons. After Erm is turned off, the second stripe of Opa together with Ey then promote the generation of Dfr^+^ Dac^+^ Notch-off neurons, possibly acting together with spatial factors. In summary, our data showed that the Ey stage is divided into (at least) two sub-temporal stages with Ey/Erm, and Ey/Opa as TTFs, respectively, and these different combination of TTFs determine different neural fates. (Figure 3L)

### Scro is required to promote Slp expression to the threshold level for temporal transition

Based on our scRNA-seq data, scro mRNA is expressed in NBs starting at a similar time as Slp 1 and 2. The *scro* gene encodes a NK-2 homeobox transcription factor, the expression of which in the medulla has been indicated by its knock-in mutant alleles in a recent study ^71, 72^.

To study the function of Scro in the temporal cascade, we generated *scro* RNAi clones, and observed that in such clones, Slp expression was greatly reduced (Figure 4A,A’), while Opa and Ey expression was expanded into older NBs, and D expression was lost (Figure 4B-D’). This set of data suggest that Scro promotes the transition from the Ey stage to the Slp stage by activating Slp expression. Since Slp is required to repress Ey and Opa, as well as to activate D, the expansion of Opa and Ey as well as the absence of D expression in *scro* RNAi clones might be due to the weak Slp expression caused by loss of Scro, but a direct role for Scro also cannot be excluded (Figure 4J). Next, we tested Scro’s effect on the neuron fate generated in the Slp temporal window. Sox102F is a transcription factor expressed in subsets of neuronal progeny of the Slp stage and D stage NBs (Figure S4A), and it is lost in *slp* mutant clones^40^. In *scro* RNAi clones, Sox102F^+^ neurons were also largely lost, showing that Scro is required for the neuron fate generated in the Slp stage (Figure 4E,E’). Thus our data suggest that Scro is required to activate Slp expression to the full level that allows the actual transition to the Slp stage to occur (Figure 4J), because with loss of Scro, although a weak level of Slp is still expressed in NBs, it is not sufficient to specify the correct neural type, or promote temporal progression of the cascade.

**Figure 4.**
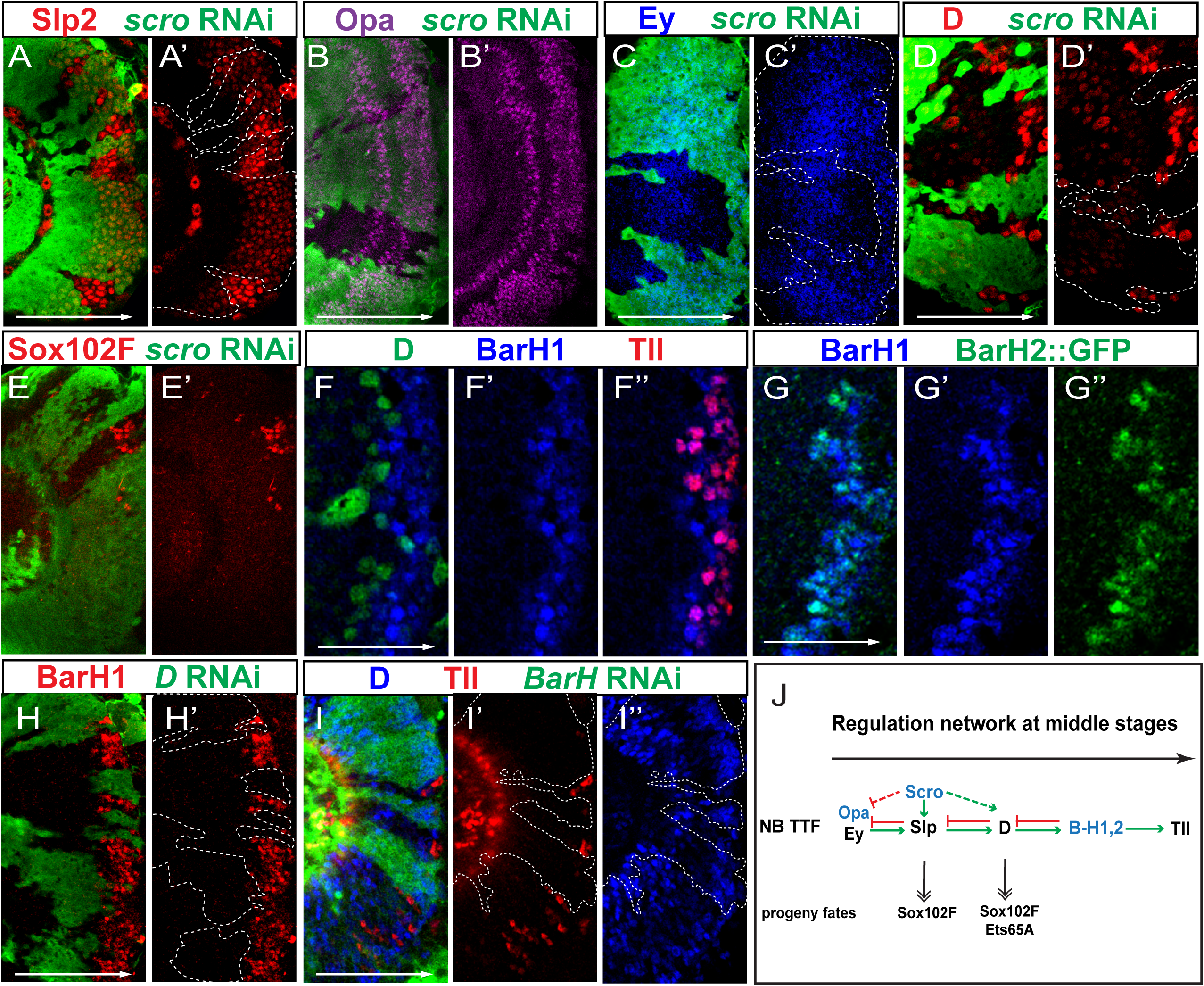
Scro and BarH proteins are among late TTFs. (A-A’) In *scro-RNAi* (BDSC 33890) clones marked by GFP (green), Slp expression (red) is much weaker compared to outside of the clones (9 out of 9 clones). (B-B’) In *scro-RNAi* (BDSC 33890) clones marked by GFP (green), Opa expression (magenta) is expanded into older NBs (8 out of 8 clones). (C-D’) In *scro-RNAi* (BDSC 33890) clones marked by GFP (green), Ey expression (blue) is expanded into older NBs, and D expression (red) is lost (8 out of 8 clones). The D antibody we used has unspecific cross-reactivity with other TFs in young neuroblasts (weak staining), but it specifically recognizes D at the D stage, which is demonstrated in Figure S2F. (E,E’) In *scro-RNAi* (BDSC 29387) clones marked by GFP (green), Sox102F (red) expressing neurons are not generated (15 out of 17 clones show a complete loss). (F-F’’) The expression pattern of D (green), BarH1 (blue) and Tll (red) in NBs. (G-G’’) The expression of BarH1 (blue) and BarH2::GFP (green) in NBs. (H-H’) In *D-RNAi* clones marked by GFP in green, BarH1 expression (red) is lost. (I-I’’) In *BarH1* and *BarH2* double RNAi clones marked by GFP (green), Tll expression (red) is lost, and D expression (blue) is expanded (in 15 out of 15 clones). (J) A schematic showing the regulatory network among late TTFs: Scro, Slp, D, BarH, Tll, and the neuron fates generated at each stage.

### BarH genes mark the temporal stage between the D stage and the Tll stage

*BarH1* and *BarH2* are two homologous genes encoding homeobox transcription factors, best known for their function in the development of the eye^73, 74^. In the medulla, consistent with our scRNA-seq data, both genes are expressed in medulla NBs between the D stage and the Tll stage. D, BarH proteins, and Tll are expressed in three consecutive stripes with BarH proteins in the middle (Figure 4F-F’’, Figure S4B-B’’). While D expression and Tll expression do not overlap with each other, the expression of BarH1 and BarH2 overlaps with both D and Tll. BarH1 and BarH2 are initiated at almost the same time, and their stripes in NBs are much overlapped, but the highest level of expression of the two TFs is achieved at different times in different cells (Figure 4G-G’’). The inequivalence of their expression is amplified in neurons that are born at the BarH stage, as neurons expressing only BarH1 or BarH2 are observed inside the medulla (Figure S4C-C’’).

If BarH1 and BarH2 are TTFs between D and Tll, they might be activated by D, and be responsible for terminating D expression and activating Tll expression. We used *ay*G4 to drive *D-*RNAi, and observed loss of BarH1 expression in *D*-RNAi clones (Figure 4H,H’), suggesting that D is required for BarH1 activation. To test whether BarH1 and BarH2 are required for the transitions from the D stage to the Tll stage, we generated clones in which RNAi of both BarH genes were induced. Inside such RNAi clones, Tll was lost while D was expanded to the oldest NBs (Figure 4I-I’’). Next, we tested whether BarH1 and BarH2 are individually required in the transitions. With loss of BarH1 or BarH2 alone, normal Tll expression was observed in NBs (Figure S4D, F), suggesting that BarH1 and BarH2 act redundantly to activate the next TTF Tll (Figure 4J).

### Gcm is the final TTF required for the switch to gliogenesis and cell cycle exit

Previously it was thought that Tll is the last TTF expressed in the oldest medulla neuroblasts that produce glia, but whether Tll indeed plays a role in gliogenesis hasn’t been examined^18^. Our scRNA-seq data suggest that there is another temporal stage after the Tll stage marked by the expression of Gcm and Dacapo (Dap) (Figure 1G, FigureS5A). Gcm, a zinc finger transcription factor, was shown to be essential for glial fate determination in the embryonic nervous system and larval visual system^75–78^. Then another study showed that Gcm is expressed in a group of precursors located at the border between the optic lobe and the central brain and required to generate medulla neuropil glia (mng)^41^. Our scRNA-seq data suggest that these precursors are the final stage of medulla neuroblasts, rather than a separate group of dedicated glial precursors.

A Gcm::GFP-BAC line was used to examine the protein expression pattern of Gcm, and it was confirmed that Gcm protein is expressed after Tll in the oldest medulla NBs (Figure 5A-A’’ and Figure S5B-B’’). Gcm expression has a significant overlap with Tll, but in a more restricted stripe closer to the central brain. A high level of Gcm expression is often observed in neuroblasts with a reduced level of Tll. Gcm-expressing progeny later activate Repo expression, as suggested by the co-expression of Gcm::GFP and Repo in the migrating mng generated by the oldest medulla NBs (Figure S5C-C’’). As mng migrate towards the medulla neuropil, Gcm expression reduces, while the expression of Gcm2, a homolog of Gcm, increases (Figure S5D-D’’). Gcm2 is not expressed in the neuroblasts^41^ (Figure S5D-D’’). Although some Gcm-expressing NBs and newly generated mng still transiently retain a low level of Tll, most Tll-expressing progeny do not express Gcm or Repo, instead, some of them express Dac (Figure S5E-E’’), suggesting that Tll is a TTF activated before Gcm, and that Tll stage NBs produce Tll^+^ neurons including some expressing both Tll and Dac. These neurons appear to only express Tll transiently, and Tll is turned off in mature neurons.

**Figure 5.**
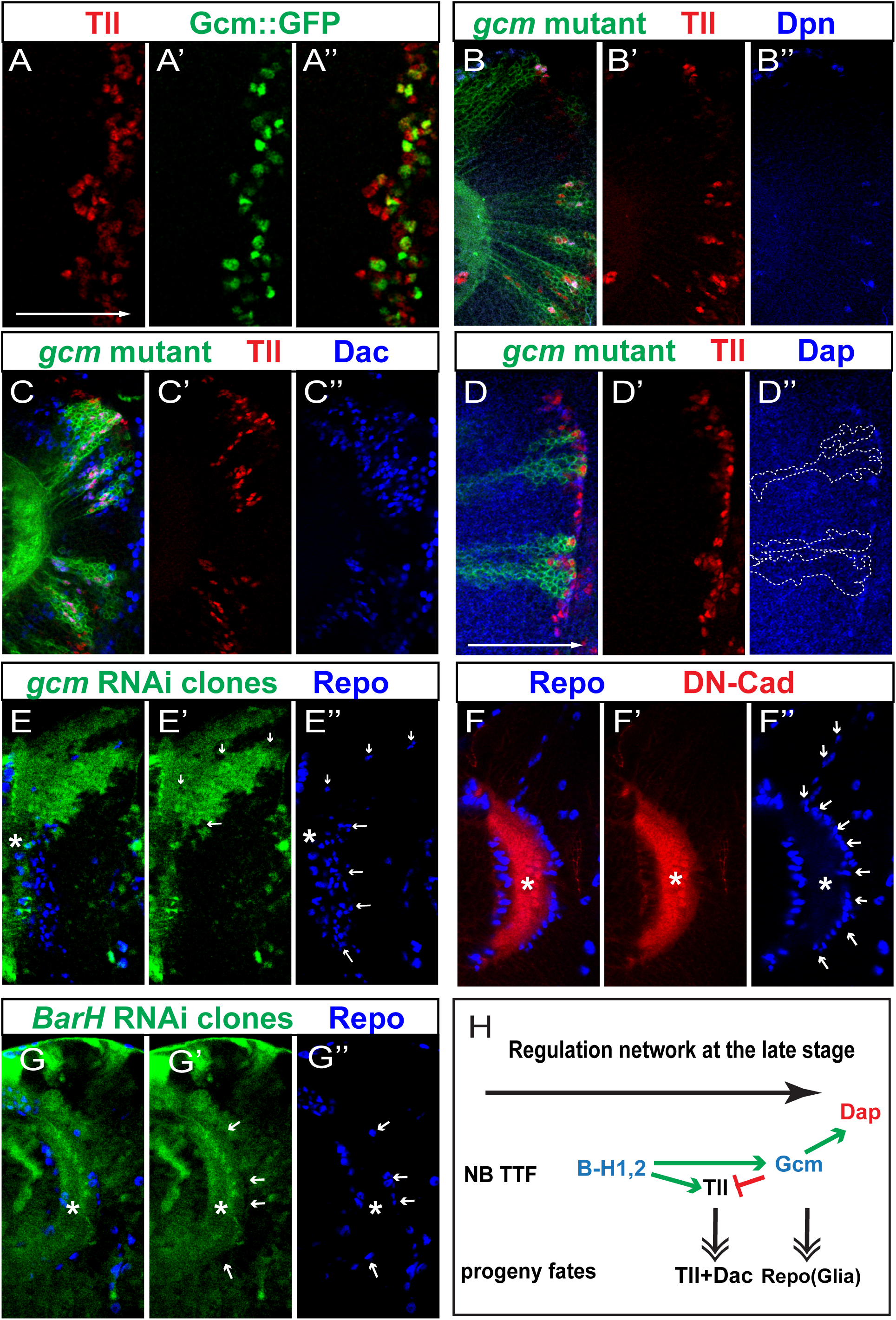
Gcm is the TTF that terminates the temporal cascade. (A-A’’) The expression of Tll (red) and Gcm::GFP (green) in NBs. (B-B’’) In *gcm* mutant clones (marked by GFP in green), ectopic NBs marked by Dpn (blue) and Tll (red) are present in a deep progeny focal plane, along with ectopic Tll^+^ progeny surrounding the ectopic NBs (in 17 out of 17 clones). (C-C’’) In *gcm* mutant clones (green), Tll (red) and Dac (blue) double positive cells are increased (in 15 out of 15 clones). (D-D’’) In *gcm* mutant clones (green), Dap (blue) is lost. The co-expression of Tll (red) and Dap is only present outside the clones (in 6 out of 6 brains). (E-E’’) In *gcm* RNAi clones (marked by GFP in green), glia marked by Repo (blue) are mostly lost (in 12 out of 12 clones). The asterisk indicates the center of the neuropil, and the arrows are pointing at the glia. Note the glia don’t have GFP expression. (F-F’’) In wild-type brains, mng marked by Repo (blue) migrate towards the neuropil marked by DN-Cad (red), and are aligned continuously around the neuropil surface. The asterisk indicates the center of the neuropil, and the arrows are pointing at the glia. (G-G’’) In *BarH1* and *BarH2* double RNAi clones marked by GFP (green), mng marked by Repo (blue) is greatly lost (in 6 out of 6 clones). The asterisk indicates the center of the neuropil, and the arrows are pointing at the glia. The remaining glia don’t have GFP expression. (H) A schematic showing the regulatory network of Gcm and its relating TTFs: BarH, Tll, and the progeny fates generated at the Gcm stage and the Tll stage.

Thus, Gcm is expressed at the final stage when NBs transit from neurogenesis to gliogenesis and subsequently exit the cell cycle. Dap, a cell cycle inhibitor orthologous to vertebrate Cdkn1a/P21, is expressed in a similar pattern as Gcm (Figure 1G, FigureS5A). Tll is expressed in a slightly earlier stage, and loss of Tll does not affect the cell cycle exit or glia production (Figure S5F-F’’). Thus, Gcm might be the critical regulator that functions in promoting gliogenesis and cell cycle exit.

We examined the function of Gcm by generating *gcm* mutant clones. In a wild-type brain, cells expressing both Dpn and Tll are not present deep inside the medulla where only neurons and glia are located. However, with loss of *gcm*, ectopic Tll^+^ Dpn^+^ cells are observed at deeper planes (Figure 5B-B’’). Those cells should be the Tll^+^ neuroblasts unable to transit into the glia producing and cell cycle exiting mode, and instead, they remain at the neuron producing mode and keep producing supernumerary Tll^+^ Dac^+^ neurons (Figure 5C-C’’). In *gcm* RNAi clones, we also observed excessive Tll^+^ Dac^+^ neurons (Figure S5G-G’’). These results suggest that Gcm is required to repress Tll, and end the temporal progression. The extended NB proliferation could be due to the loss of Dap expression in *gcm* mutant clones (Figure 5D-D’’). Confirming Gcm’s known role in gliogenesis, we never observed Repo^+^ glia around the medulla neuropil inside *gcm* mutant clones or *gcm* RNAi clones (Figure 5E-E’’). To determine if Gcm is sufficient to promote gliogenesis and terminate neuroblast proliferation, we tested the effect of misexpression of Gcm in younger NBs. Clones mis-expressing Gcm were small, and composed mostly of neuroblasts marked by Dpn and glia marked by Repo, suggesting that Gcm is sufficient to promote gliogenesis, but may cooperate with other factors to promote cell cycle exit (Figure S5H-H’’). In summary, this set of data support that Gcm is the final TTF required for the switch to gliogenesis and for ending the temporal cascade by activating Dap.

As Tll is not required for glia production suggesting that Tll is not required to activate Gcm, there should be another TTF directly upstream of Gcm and required for Gcm expression. Therefore, we tested if *BarH* genes are the upstream TTFs. Because of the lack of Gcm antibody, we used Repo expression as an indicator for the successful progression towards the final temporal stage. While mng produced by the oldest NBs line up around the medulla neuropil continuously in a wild-type brain (Figure 5F-F’’), in *BarH1* and *BarH2* double RNAi clones, mng was lost, suggesting that BarH1 and BarH2 are upstream of Gcm in the temporal cascade (Figure 5G-G’’). However, mng production was normal with individual RNAi of *BarH1* or *BarH2*, suggesting that BarH1 and BarH2 act redundantly to promote the temporal cascade towards the final stage (Figure S4E, G). In summary, BarH1 and BarH2 are required to activate both Tll and Gcm, but with different kinetics, and Gcm is then required to promote gliogenesis and cell cycle exit by repressing Tll and activating Dap (Figure 5H).

### TFs functioning along the differentiation axis participate in the temporal cascade regulation

Our scRNA-seq analysis enabled us to identify a fairly complete temporal cascade from start to end. An important question concerning the regulation of the temporal cascade is why the progression of the TTF cascade is only present in NBs, but not in neurons. Our hypothesis is that factors differently expressed between NBs and neurons might contribute to limit the sequential activation of TTFs to NBs. To test this hypothesis, we screened through some TFs that are not expressed in a TTF manner but rather differentially expressed along the differentiation axis (NBs -> GMCs -> neurons), and we found that two genes, *longitudinals lacking* (*lola*) and *nervous fingers 1* (*nerfin-1*), participate in the temporal cascade regulation.

The gene *lola* encodes about 20 isoforms of transcription factors belonging to a Broad-complex, Tramtrack and Bric-à-brac/poxvirus and zinc finger (BTB/POZ) family of proteins. The isoforms have a common BTB domain and different Zinc fingers that give each isoform different DNA binding specificities ^79^. The 20 isoforms show different expression patterns in the medulla. For example, Lola-F (isoform nomenclature as in Ref^79^) is expressed in all NBs, all GMCs and newly born neurons, but is down-regulated quickly to absence as neurons mature (Figure 6A-A’’, Figure S6A-A’’). Lola-N is expressed mostly in mature neurons^80^. A Lola-T::GFP-BAC line showed weak Lola-T::GFP expression in a similar pattern as that of Lola-F, although with a much earlier activation starting from NE (Figure 6B,B’). A Lola-K::GFP-BAC line showed that Lola-K is expressed at a high level in mature neurons and NBs, but is not detected in GMCs (Figure 6C,C’). Therefore, different isoforms of Lola could have diverse functions in NBs, GMCs and neurons. Overall, this change in Lola isoform composition happens along the differentiation axis. For example, only NBs express both Lola-F and Lola-K, which might act together to regulate NB-specific processes. Using *optix*G4 which is expressed in the mOPC ^65^, or *ayG4* to drive *lola* RNAi that eliminates all isoforms of Lola, we observed a great expansion of Hth expression in NBs, a slight expansion of the first stripe of Opa, and delays in Ey expression, the second stripe of Opa, and Slp expression to increasing extents in NBs (Figure 6D-G’’, Figure S6B-D’’). The first stripe of Opa was only slightly expanded, but the second stripe of Opa is further delayed as the gap between the two stripes of Opa became much larger; the delay of Ey expression was very severe and the Slp expression was often absent. This is as if the temporal progression grows slower and slower (Figure 6P). With *lola* RNAi driven by *optixG4* or *ayG4*, we also observed a great reduction of Runt neurons and Kn neurons, suggesting that Lola proteins are also required for the neuronal fate specification (Figure 6E and data not shown). This set of data suggest that Lola inhibits Hth expression and facilitates the normal temporal progression, and Lola might be a speed modulator of the temporal cascade in NBs (Figure 6P).

**Figure 6.**
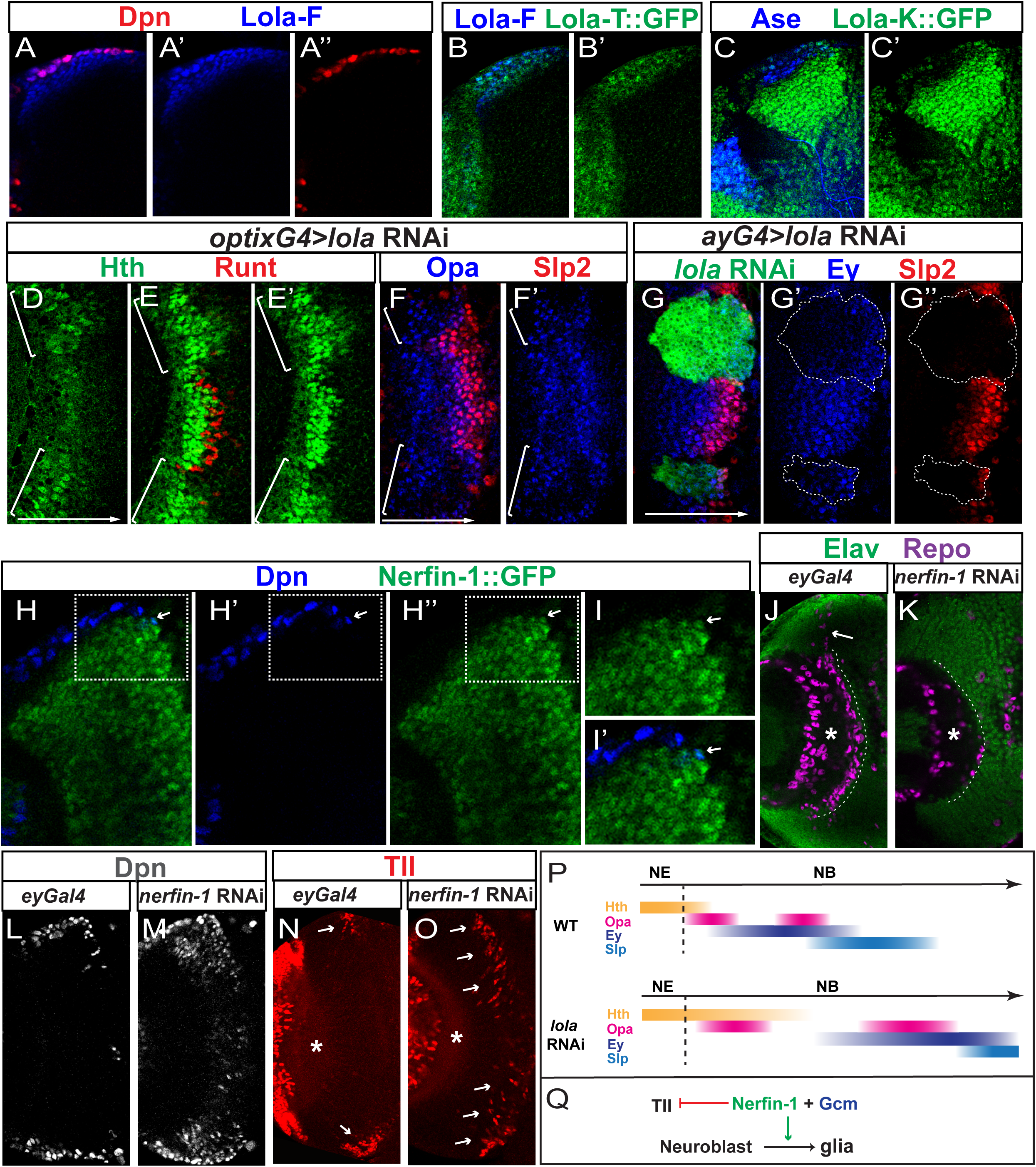
Lola and Nerfin-1 regulate the progression and termination of the temporal cascade, respectively. The *lola* RNAi line used in this figure is BDSC35721. (A-A’’) The expression of Dpn (red) and Lola-F (blue) in a cross-sectional view. (B,B’) The expression of Lola-T::GFP (green) and Lola-F (blue) in a cross-sectional view. (C,C’) The expression of Lola-K::GFP (green) and Ase (blue) that marks both NBs and GMCs in a cross-sectional view. (D-F’) *lola* RNAi is driven by *optixGal4* in mOPC indicated by white brackets. (D-E’) With *lola* RNAi, Hth (green) is expanded into older NBs (D) and later-born progeny (E,E’), while Runt neurons (red) are mostly lost (E) (in 5 out of 5 brains). (F-F’) With *lola* RNAi, the expression of the first stripe of Opa (blue) is slightly expanded, while the expression of the second stripe of Opa is greatly delayed (in 7 out of 7 brains). (G-G’’) In *lola* RNAi clones, the expression of Ey (blue) and Slp2 (red) are greatly delayed (in 16 out of 16 clones). (H-H’’) The expression of Dpn (blue) and Nerfin-1::GFP (green) in a cross-sectional view. The right most NB is the oldest NB that turns on Nerfin-1 (arrow). (I-I’) A zoomed-in image of the outlined region in panel H. (J,K) In deep progeny focal planes, glia are marked by Repo (magenta), and neurons are marked by Elav (green). The asterisk indicates the center of the medulla neuropil, and the white dashed line indicates the position where mng should be aligned. (J) In *ey*Gal4 control brains (n=5), mng continuously align the medulla neuropil, and the migrating mng stream is indicated by a white arrow. (K) When *Nerfin-1* RNAi is driven by *eyGal4*, only scattered mng are observed (in 5 out of 5 brains). (L) At a focal plane slightly deeper than the surface focal plane, most NBs marked by Dpn in an *eyGal4* control brain are located at the surface of the medulla. (M) When *Nerfin-1* RNAi is driven by *eyGal4*, many ectopic NBs marked by Dpn are present inside the medulla (in 5 out of 5 brains). (N) At a deep progeny focal plane, Tll^+^ (red) progeny (white arrows) are only generated around the surface NBs, while no Tll+ cells are observed in the middle of the brain. (O) At a comparably deep progeny focal plane, when *Nerfin-1* RNAi is driven by *eyGal4*, Tll (red) expressing progeny continue to be produced throughout the brain (in 6 out of 6 brains). (P) A schematic showing the function of Lola in regulating the temporal cascade. Loss of Lola causes slowing down of the temporal progression. (Q) Nerfin-1 is required for the transition from the Tll stage to gliogenesis.

Nerfin-1 is a zinc-finger transcription factor expressed in postmitotic neurons and is required for maintaining their differentiated status^81, 82^. According to our scRNA-seq data, Nerfin-1 transcripts are present in the oldest NBs similar to Gcm transcripts (Figure 1G,H). Using a Nerfin-1::GFP-BAC line, we showed that Nerfin-1 protein is expressed mostly in maturing neurons as previously reported (Figure 6H). However, the co-expression of Dpn and Nerfin-1 can be observed in the nuclei of the oldest NBs (Figure 6H-I’). Nerfin-1 expression persists in the new-born glia generated by the oldest NBs for a short time and is lost as the glia mature and migrate, suggested by the fact that Repo seldom co-expresses with Nerfin-1 in the mng glia located around the neuropil (data not shown). Next, we tested whether Nerfin-1 is required for the cell cycle exit and glia production. We used eyG4 to drive *nerfin-1* RNAi, and showed that Nerfin-1 is another critical regulator for the final-stage NBs. In a wild type control brain, mng produced from the oldest NBs line up the medulla neuropil continuously (Figure 6J). In contrast, with loss of Nerfin-1, only a few scattered mng are observed around the medulla neuropil (Figure 6K)(the reduction is highly significant: p=1.15X10^-10^ by t-test, n=5 brains each). This suggests that the transition to gliogenesis is affected by the loss of Nerfin-1, and it is possible that NBs could be stuck at the previous Tll stage. With loss of Nerfin-1, we also observed numerous ectopic Dpn^+^ neuroblasts inside the medulla (Figure 6M), consistent with previous reports showing that loss of Nerfin-1 causes de-differentiation of neurons back into neuroblasts^81, 82^. However, if the oldest NBs fail to exit the cell cycle, they can also be among these ectopic neuroblasts. To test this possibility, we examined Tll expression. Tll is transiently expressed in the newly-born progeny from Tll^+^ NBs, and will be lost soon after. In a wild type brain at a deeper focal plane, NBs are only observed at the surface, and Tll is observed in old NBs and their newly-born progeny just below the surface (Figure 6N arrows). The lineages in the middle of the brain have been completed (NBs have finished generating mng and exited the cell cycle), and thus no NB or Tll^+^ progeny is observed (Figure 6N). However, with loss of Nerfin-1, Tll^+^ progeny continue to be produced throughout the brain (Figure 6O). This set of data suggest that Nerfin-1 is also required for the transition from the Tll stage to gliogenesis and for exiting the cell cycle. Thus Nerfin-1, which is required in neurons to maintain their differentiation status, is turned on in the oldest NBs to promote gliogeneis and put a stop on the temporal progression (Figure 6Q).

Together, the requirement of Lola and Nerfin-1 in the normal cascade progression and termination suggests that genes that are differentially expressed and functioning along the differentiation axis also contribute to the temporal cascade regulation.

## Discussion

### ScRNA-Seq is a powerful tool to study temporal patterning

Our scRNA-Seq analysis revealed the temporal progression of transcriptional profiles as medulla neuroblasts age at single-cell resolution. We discovered candidates of critical temporal patterning regulators including eight novel TTFs, as well as TFs such as Lola and Nerfin-1, which are expressed along the differentiation axis but also involved in the temporal patterning process. Further experimental validation of novel TTFs and other crucial regulators confirmed the accuracy of our high-resolution data, supporting that scRNA-seq is a powerful tool to study the highly dynamic temporal patterning process. Our analysis and further experimental investigation revealed a comprehensive temporal cascade in *Drosophila* medulla neuroblasts: Hth + SoxN + dmrt99B -> Opa -> Ey+Erm -> Ey+Opa -> Slp+Scro -> D -> BarH1&2->Tll, Gcm (Figure 7B), and also illustrated several principles that might be conserved during the temporal patterning of neural stem cells.

**Figure 7.**
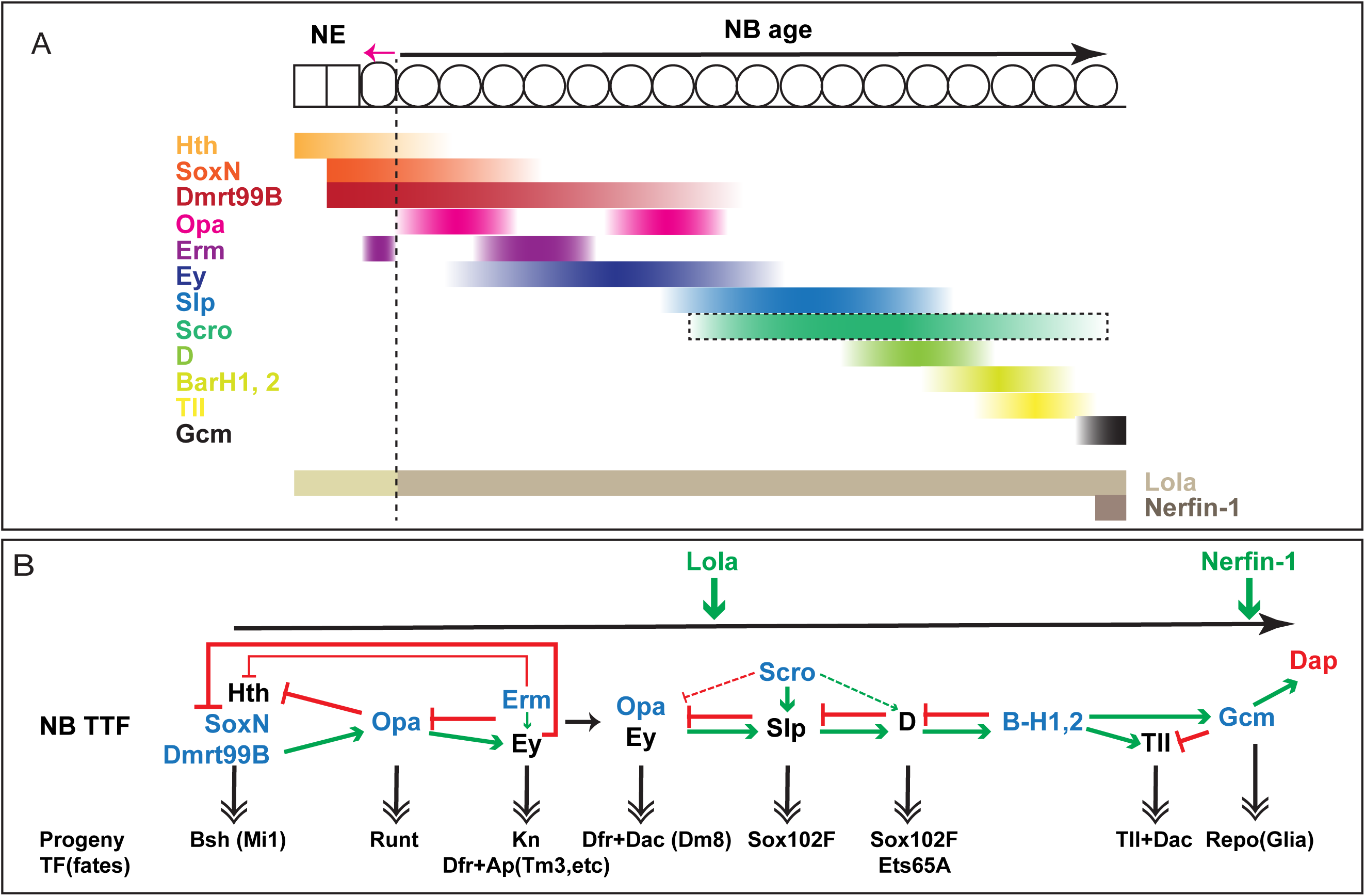
A schematic model summarizing the medulla TTF cascade and its regulation. (A) A schematic drawing showing the relative expression patterns of the medulla TTFs, Lola and Nerfin-1. The expression of Scro is indicated only by its transcriptional pattern. The expression of Hth, SoxN and Dmrt99B all start in NE, Em has a stripe in the transition zone from NE to NB, while other TTFs are initiated in NBs. Different isoform compositions of Lola are indicated by different colors. The number of NE and NB cells do not indicate the actual number of cell cycles they go through. (B) A schematic model summarizing the regulation networks of the medulla TTF cascade. Known TTFs are in black, and new TTFs identified in this study are in blue. Extensive cross-regulations were identified between these TTFs, that generally follow the rule that a TTF is required to activate the next TTF (green arrows) and repress the previous TTF (red flat-headed arrows), but with a few important exceptions. This TTF cascade controls the sequential generation of different neural types by regulating the expression of neuronal transcription factors, and examples of neural types were also indicated. Lola proteins modulate the speed of temporal progression of the NB TTF cascade. At the final stage, Gcm and Nerfin-1 promote gliogenesis and the cell-cycle exit to end the temporal progression.

### Initiation of the TTF cascade

Before this study, Hth was proposed to be the only TTF at play during the earliest temporal stage. Hth is expressed in the neuroepithelium and the youngest NBs. It is necessary for the generation of Bsh neurons, but is required neither for the NE to NB transition, nor for the further temporal cascade progression. Loss of Ey also does not affect the termination of Hth ^18, 19^. These data suggested missing links between Hth and the later TTF cascade. Here we identified several novel TTFs that linked the whole cascade together. Two novel TTFs that started their expression in the NE, SoxN and Dmrt99B, are also required for the first temporal fate (Bsh neurons), and Dmrt99B is required for the timely activation of Opa in the youngest NBs. Opa is then required to activate Ey and repress Hth (Figure 7A,B). Interestingly, the three TTFs inherited from NE maintain their expression for different durations in NBs, as Hth is repressed by Opa and Erm, SoxN is repressed by Ey, while Dmrt99B expression extends until the Slp stage (Figure7A, B). Whether this differential down-regulation is significant for temporal patterning is currently unknown. However, it is worth noting that the expression of mammalian orthologs of Dmrt99B, Dmrt3 and Dmrta1, also starts in symmetrically dividing early cortical progenitors (NE), and decreases gradually in asymmetrical dividing cortical progenitors due to the direct suppression by FoxG1, the mammalian ortholog of Slp1/2 ^83, 84^. Given the essential role of Dmrt99B in initiating temporal patterning in medulla neuroblast, it will be interesting to investigate whether its mammalian orthologs play conserved roles in the temporal progression of cortical progenitors.

### A broad temporal stage can be divided into sub-temporal stages by combinations of TTFs

At certain temporal stages, not just one TTF, but combinations of TTFs determine the progeny fates. This is well-illustrated in the Ey stage. The first stripe of Opa is required to initiate Ey expression in NBs, and Erm (as shown by scRNA-seq and Erm::V5) is turned on at a similar time as Ey. Due to the lack of Erm antibody, we haven’t examined how Erm expression is regulated, but judging from the expression pattern, it is possible that Erm activation might also require Opa. Erm then transiently represses Opa until itself is tuned off possibly due to the loss of Opa, forming a negative feedback loop. Negative feedback loops with time-delay can generate striped expression patterns. After Erm is turned off, Opa is turned back on. At the same time Slp has been gradually activated by Ey and Scro, and when it reaches a certain level, it will repress Opa and Ey to end the Ey stage. Thus, cross-regulations among TTFs divide the Ey stage into (at least) two (sub-)temporal stages determined by the co-expression of Ey and Erm, or Ey and Opa. We showed that different neural types are generated in these two sub-temporal stages, and the first set of neurons require both Ey and Erm, while the second set of neurons require both Ey and Opa (Figure 7A,B). Interestingly to note, the mammalian ortholog of Erm, Fezf2, is also expressed in cortical progenitors and play important roles in cortical neuron specification ^34, 35, 85^.

### Termination of the TTF cascade

Previously it was thought that Tll stage NBs switch to gliogenesis and then exit the cell cycle, but whether Tll indeed plays a role in these processes has not been studied. Here our scRNA-Seq data suggested another final temporal stage marked by the expression of Gcm and Dap. Further we showed that BarH1 and BarH2 are required to activate both Tll and Gcm, but with different kinetics. Tll is activated first, and when Gcm is activated, Gcm represses Tll. We showed that Gcm but not Tll is required for the NBs to switch to gliogenesis and exit the cell cycle (Figure 7 A,B). Gcm is well-known for its role in gliogenesis, but here we show that it is also required to activate Dap expression in neuroblasts to promote cell cycle exit and end the temporal progression. In vertebrate retina, scRNA-seq analysis of retinal progenitor cells identified NFI factors as required for both late-born cell fates including Müller glia and for exiting the cell cycle^28^. Since neural progenitors often switch to produce glia at the end of the lineage, it might be a general mechanism that factors required for the switch to gliogenesis are also required for the mitotic exit to end the temporal progression.

### Complex cross-regulations among TTFs form temporal gene networks

The model for the cross-regulations between medulla TTFs is that each TTF activates the next TTF and inhibits the previous TTF from the Ey stage to the end of the cascade, exhibiting a simple combination of feedforward activation and feedback repression. However, based on the experimental evidence we produced as well as inferred from the scRNA-seq data, the cross-regulations among TTFs are more complex. One TTF is not necessarily repressed by the very next TTF, or activated by the exactly previous TTF. Hth is repressed by Opa and Erm. SoxN is repressed by Ey, while Dmrt99B is likely to be repressed by Slp or later TTFs. Tll is activated just before Gcm, however, Tll is not required for Gcm’s activation. The complexity of their cross-regulation is a way to increase the number of combinations of TTFs in aging NBs, thereby increase the number of possible neuron fates determined along the temporal progression. However, the overall trend that early TTFs activate late TTFs, and late TTFs repress early TTFs remains valid.

### Genes functioning in the differentiation axis regulate the TTF cascade

A novel mechanism for the regulation of temporal patterning is discovered in our study: TFs differentially expressed and functioning along the differentiation axis also contribute to the regulation of the TTF cascade (Figure 6P,Q). There are two temporal axes in a NB lineage, one is the TTF cascade progression in NBs, and the other is the gradual differentiation of the progeny losing the self-renewal ability (NBs to GMCs to neurons). Whether the two axes interact with each other was not clear. In this study we investigated the role of two TFs, Lola and Nerfin-1, that are regulated and functioning along the differentiation axis, in the TTF cascade progression. Lola proteins belong to a BTB/POZ family of proteins which have been shown to be involved in chromatin remodeling and organization^86^. Certain isoforms of Lola are expressed in all NBs, e.g. Lola-F is activated one cell cycle earlier than Opa. We show that Lola proteins functions as a speed modulator of the temporal cascade progression. It represses the expression of Hth, facilitates the activation of Opa and the following TTFs to different extents, and guarantees a quick transition from the NE gene network to the NB gene network. Interestingly, the vertebrate ortholog of *lola*, Zbtb20, was also found to modulate the sequential generation of different neural types in cortical progenitors^87^. Loss of Zbtb20 causes the temporal transitions to be delayed further and further, very similar to the loss of *lola* phenotype in our system. Thus, it is possible that lola / Zbtb20 play conserved roles in temporal patterning of neural progenitors.

The expression of Nerfin-1 is also differentiation-axis-dependent. It is observable mostly in maturing neurons, and is required to prevent neurons from dedifferentiation^81, 82^. However, this TF responsible for maintaining the differentiation status of neurons, is turned on in the final stage NBs, where it functions to promote gliogenesis and help terminate the temporal cascade on time. The fast exit of cell cycle at the final stage might be accomplished because self-renewal repressors that usually function in GMCs and neurons, such as Prospero^18^ and Nerfin-1, gather and cooperate in the oldest NBs. Since Gcm is also required to promote gliogenesis and cell cycle exit, Gcm and Nerfin-1 might regulate each other’s expression or cooperate with each other in this process.

In summary, the entire life of a medulla neuroblast from the beginning to the end was revealed in this study. Our comprehensive study of the medulla neuroblast temporal cascade illustrated mechanisms that might be conserved in the temporal patterning of neural stem cells. The single cell RNA-sequencing data provide a plethora of information that allow further exploration of the mechanisms of temporal patterning.

## Methods

### Fly lines and crosses

#### Construction of fly lines

To construct the stock for labeling of medulla neuroblasts for FACS sorting and scRNA-seq, *SoxNGal4* (GMR41H10Gal4)^88^ was recombined with *UAS-RedStinger* (BDSC 8547) on Chromosome III, and then crossed with *E(spl)mγGFP* on II ^89^, to generate the *E(spl)mγGFP* ; *SoxNGal4 UAS-RedStinger* /TM6B stock. To generate *SoxN* mutant clones, *SoxN^NC14^* mutation (BDSC 9938) was recombined onto FRT40A chromosome, to generate the *FRT40A SoxN^NC14^* stock.

#### MARCM mutant clones with FRT40A mutants

To generate MARCM mutant clones of *SoxN* mutant, *slp* mutant, *erm* mutant, or *gcm gcm2* double mutant, virgin females of *yw hsFLP UASCD8GFP; FRT40A tubGal80; tubGal4/TM6B* were crossed with males of *FRT40A SoxN^NC14^/CyO*, *FRT40A slp^S37A^* /*SM6-TM6B* (Gift from Andrew Tomlinson ^90^), *FRT40A erm^1^/CyO,GFP* (gift from Cheng-Yu Lee ^66^), or *Df(2L)200 FRT40A / Gla, Bc* (which deletes both gcm and gcm2 ^78^) respectively. The progeny were grown at 25 °C, heat shocked once at 37 °C for 40min at 1st instar larval stage, and then grown at 25 °C for three days before dissection of the wandering 3rd instar larvae.

#### MARCM mutant clones with FRT82B mutants

To generate MARCM clones of *hth* mutant or *opa* mutant, virgin females of *ywhsFLP UASCD8GFP; ; tubGal4, FRT82B tubGal80 /TM6B* were crossed with *FRT82B hth^P2^ /TM6B* flies (gifts from Richard Mann), *FRT82B opa^7^* (gift from Deborah Hursh ^63^), respectively. The progeny were grown at 25 °C, heat shocked once at 37 °C for 1hr at 1st instar larval stage, and then grown at 25 °C for three days before dissection of the wandering 3rd instar larvae.

#### Negatively marked *ey* mutant clones

Females of *yw, hs-Flp^1.22^ ;; FRT80B, eyBAC, Ubi-GFP/TM6B,Tb; ey^J5.71^* were crossed to males with genotype *hs-Flp^1.22^;; FRT80B; ey^J5.71^/ In(4)ci^D^* (Ref ^18^). The progeny were grown at 25 °C, heat shocked once at 37 °C for 1hr at 1st instar larval stage, and then grown at 25 °C for three days before dissection of the wandering 3rd instar larvae. Clones in larvae that lacked both GFP fluorescence and staining with an anti-Ey antibody were further analyzed.

#### RNAi clones

RNAi lines used in this study include: *UAS-eyRNAi* (BDSC 32486), *UAS-dmrt99B-RNAi* (BDSC 31982), *UAS-opaRNAi* (VDRC 101531), *UAS-erm^RNAi^* (BDSC 50661), *UAS-scro^RNAi^* lines (BDSC 29387, and BDSC 33890 showed the same phenotypes), *UAS-D^RNAi^* (VDRC 107194), *UAS-BarH1^RNAi^* (VDRC 104681), *UAS-BarH2^RNAi^* (VDRC 11570), *UAS-tll-miRNA* (gift from Tzumin Lee ^91^), *UAS-gcm^RNAi^* (VDRC 110539, VDRC 2961 showed the same phenotypes), *UAS-lola^RNAi^* lines (BDSC 35721, BDSC 26714, VDRC 101925, all showed similar phenotypes), *UAS-nerfin-1^RNAi^* (VDRC 101631), *UAS-oaz^RNAi^* (VDRC 39214, VDRC 107061), *UAS-hbn^RNAi^* (VDRC 103979), *UAS-sba^RNAi^* (vdrc 101314), and *UAS-scrt^RNAi^* (VDRC 105201). To generate RNAi clones, virgin females of *yw hs FLP; act>y+>Gal4 UAS GFP / CyO; UASDCR2/TM6B* were crossed with males of each of the RNAi lines. The progeny were grown at 25 °C, heat shocked once at 37 °C for 8 min at 1^st^ instar larval stage, and transferred to 29 °C for three days before dissection of the wandering 3^rd^ instar larvae.

#### Region-specific RNAi

Alternatively, region-specific Gal4s combined with UAS-DCR2 were used to drive RNAi. Virgin females of *UAS-Dcr2;Dpn-LacZ/CyO; VsxGal4/TM6B* ^65^ were crossed with males of *UAS-opaRNAi* (VDRC 101531). Virgin females of *UASDCR2; optixGal4/CyO* ^65^ were crossed with *UAS-lola^RNAi^* lines (BDSC 26714, BDSC 35721). The progeny were grown at 25 °C until 1^st^ instar larval stage and transferred to 29 °C for three days before dissection of the wandering 3^rd^ instar larvae.

#### GFP-BAC and other reporter lines

Additional lines used in this study include *Dmrt99B::GFP* (BDSC 81280), *Erm::V5* (gift from Cheng-Yu Lee ^69^), *ap*^rK568^-*lacZ* ^92^, *B-H2::GFP* (BDSC 67734), *Gcm::GFP* (BDSC 38647), *gcm-LacZ* (*P{PZ}gcm^rA87^/CyO)* (BDSC 5445), *Gcm2::GFP* (BDSC 38646), *lola-T::GFP* (flybase name: *lola.GR-GFP*) (BDSC: 38661), *lola-K::GFP* (flybase name: *lola.I-GFP*) (BDSC: 38662), and *Nerfin-1::GFP* (BDSC 67385).

### Dissociation and FACS sorting of medulla neuroblasts

For each scRNA-seq experiment, 120 third instar larvae of the genotype *E(spl)mγGFP* ; *SoxNGal4 UAS-RedStinger* /TM6B were washed with PBS twice, and with 70% ethanal for 1 min, and with PBS once again. Each of the brains was dissected on ice in complete Schneider’s culture medium (Schneider’s Insect medium, supplemented with 10% fetal bovine serum, 2% Pen/Strep and 0.02 mg/mL insulin), and was directly transferred into a glass dish on ice containing DPBS. The dissection was completed within one hour, and then the supernatant (mainly DPBS) was replaced by 1 mL TrypLE with 1mg/mL collagenase I and 1mg/mL papain. The brains were incubated for 10 min at 30°C, with gentle shaking at 55 rpm. After removal of the dissociation solution, the brains were carefully washed with complete Schneider’s culture medium once and with DPBS twice. The brains were disrupted in 1.4 ml of DPBS with 0.04% BSA by manual pipetting using a P1000, and then 0.4 ml of DPBS with 0.04% BSA was added to make a total volume of 1.8 mL. The cell suspension was filtered through the cell strainer cap into a 5mL BDFalcon FACS tube. FACS sorting was done immediately after on BD FACS ARIA II with gentle settings (85 μm nozzle and low pressure of 20 psi). Trypan blue was added before sorting. Among the singlet live cells, GFP and RFP double positive cells were selected, and sorted into DPBS with 0.04% BSA.

For immunohistochemistry of unsorted cells or sorted cells after concentration, the cell suspension was placed onto a coated (poly-D-lysine) dish for 30 minutes. After fixation with 4% formaldehyde for 10 minutes, the coverslip was washed 4 times with PBS. The primary antibodies were incubated for 2 hr, and was washed 3 times with PBST. Secondary antibodies were incubated for 30 min, and was washed 3 times with PBST. The cells were mounted in mounting medium and imaged on Zeiss confocal.

### Construction and sequencing of 10x V3 Single Cell libraries

Single-cell 3’ cDNA libraries were prepared and sequenced at the DNA Services laboratory of the Roy J. Carver Biotechnology Center at the University of Illinois at Urbana-Champaign. FACS sorted cells were immediately concentrated by centrifugation at 500g for 5 minutes, then an additional 800g for 5 minutes to a 40ul volume. This entire volume was used as input for the 10x library. The single cell suspension was converted into an individually barcoded cDNA library with the Chromium Next GEM Single-Cell 3’ single-index kit version 3 from 10X Genomics (Pleasanton, CA) following the manufacturer’s protocols. The target capture was 10k cells.

Following ds-cDNA synthesis, sequencing library compatible with the Illumina chemistry was constructed. The final library was quantitated on Qubit and the average size determined on the AATI Fragment Analyzer (Advanced Analytics, Ames, IA). The final library was diluted to 5nM concentration and further quantitated by qPCR on a Bio-Rad CFX Connect Real-Time System (Bio-Rad Laboratories, Inc. CA).

The final library was sequenced on one lane of an Illumina NovaSeq 6000 S1 flowcell (exp1) or a half lane of an Illumina NovaSeq 6000 S4 flowcell, as paired-reads with 28 cycles for read 1, 8 cycles for the index read, and 150 cycles for read 2. Basecalling and demultiplexing of raw data was done with the mkfastq command of the software Cell Ranger 3.1.0 (10x Genomics). Libraries were sequenced to a depth of 2,019,439,522 total reads (1st exp.) and 2,760,057,420 total reads (2nd exp.), corresponding to 6548 cells with a 1821 median UMI counts per cell (1st exp), and 5343 cells with 7508 median UMI counts per cell (2nd exp).

### scRNA-seq analysis

The sequencing reads were aligned to Ensembl’s BDGP6.22 using Cell Ranger (version3.0.1 for1^st^ experiment, and version 3.1.0 for 2^nd^ experiment) from 10x Genomics, and gene expression levels were counted using Cellranger “Count”. In both versions of Cell Ranger, “EmptyDrops” method^93^ was used to call cells.

All subsequent analyses were performed in R (version 4.0.3)^94^ Quality control on the count data was performed using the package Seurat (version 3.2.3) ^46^. To limit the analysis to neuroblasts, only cells expressing Dpn were analyzed. Cells were also excluded if 10% or more of their reads came from mitochondrial genes. This left 777 cells from the first experiment and 2302 cells from the second experiment.

Data from both experiments were combined into a single analysis. Batch correction was performed using the standard integration workflow in Seurat. Specifically, each dataset was first separately normalized and the top 2000 most variable features were identified. These features were then used to find integration anchors, after which the data were combined using the IntegrateData function. Finally, five outlier cells with large numbers of counts were removed. The remaining 3074 cells were scaled and centered for downstream analysis.

UMAP coordinates were calculated with the RunUMAP function using the top 10 principal components of the 2000 most variable features. Clustering was performed using the FindNeighbors and FindClusters functions with a resolution of 0.9. Cell cycle scoring was performed using the CellCycleScoring function; see Supplementary Table 1 for lists of the S genes and G2/M genes used. Gene expression levels were visualized with the FeaturePlot function. Differentially expressed genes were identified using the FindAllMarkers function with an FDR cutoff of 0.05.

Developmental trajectories were inferred using the Monocle3 (version 0.2.3.0) ^48–50^ excluding the outlying clusters 8, 11, 12, and 14. Raw counts data were imported from the integrated Seurat object into a Monocle3 cell dataset object. Preprocessing was performed with the preprocess_cds function and batch correction was performed using the Batchelor algorithm ^95^ as implemented in the align_cds function. PCA and UMAP embeddings stored in the Monocle3 object were replaced with the corresponding values calculated by Seurat and stored in the Seurat object. Cells were then clustered in Monocle3 using the cluster_cells function and the principal graph was learned using the learn_graph function with the minimal branch length option set to 5. The root node was set to be the vertex closest to cells with the highest median Hth expression. Finally, pseudotimes were calculated using the order_cells function and the trajectory was visualized using the plot_cells function. Genes with temporally patterned expression gradients were identified as those whose expression levels showed significant Spearman correlation with the inferred pseudotime at an FDR cutoff of 0.05. The top 200 genes with decreasing or increasing gradients, respectively, were analyzed for enriched Go terms for Biological Processes “GOTERM_BP_DIRECT” using the “Functional Annotation Chart” at the DAVID Bioinformatics Resources 6.8 website (https://david.ncifcrf.gov/home.jsp).

### Antibodies and Immunostaining

These antibodies are generous gifts from the fly community: Rabbit anti-SoxN (1:100) from Steven Russell ^55^; Rabbit anti-Hth (1:500) from Richard Mann; Guinea-pig anti-Run, Rabbit anti-Bsh, Rabbit anti-Slp1, Guinea-pig anti-Slp2, Rabbit anti-D, Guinea-pig anti-Tll and Rabbit anti-Sox102F (all used at 1:500) from Claude Desplan; Rabbit anti-D (1:1000) from John R. Nambu ^96^; Rat anti-Dfr (1:200) from Makoto Sato^57^, Guinea-pig anti-Kn (1:500) from Adrian Moore^97^, Guinea-pig anti-Dpn (1:500) from Chris Doe; Rabbit anti Opa (1:100) from J. Peter Gergen ^61^; Rat anti-BarH1 (1: 200) from Tiffany Cook ^98^.

Commercially available antibodies include: sheep anti-GFP (1:500, AbD Serotec, 4745-1051), Goat anti-beta-gal (1:1000, Abcam, ab12081), rabbit anti-RFP (1:1000, Abcam, ab62341), Mouse anti V5-Tag:DyLight®550 (1:200, Bio-Rad, MCA1360D550GA), Rat anti-Deadpan [11D1BC7] (1:200, Abcam, ab195173). These antibodies are provided by the Developmental Studies Hybridoma Bank (DSHB): mouse anti-eyeless (1:10), mouse anti-Pros (MR1A 1:10), mouse anti-Repo (8D12 anti-Repo 1:50), and mouse anti-Dac (mAbdac2-3, 1:20), mouse anti-Dap (NP1, 1:5), mouse anti-Lola -F (7F1-1D5, 1:20).

Secondary antibodies are from Jackson or Invitrogen. Immunostaining was done as described ^18^ with a few modifications: 3^rd^ instar Larval brains were dissected in 1XPBS, and fixed in 4% Formaldehyde for 30 minutes on ice. Brains were washed and then incubated in primary antibody solution overnight at 4°C, washed three times and incubated in secondary antibody solution overnight at 4°C, washed three times and mounted in Slowfade. Images are acquired using a Zeiss Confocal Microscope. Figures are assembled using Photoshop and Illustrator.

### Quantifications and Statistical Analysis

For each experiment, the numbers of animals or clones analyzed are included in each figure’s legends. Sample sizes are estimated based on previous experience. Clonal experiments have internal controls: we compared gene expression in and outside of the clones in the same sample, and only draw conclusions when consistent results are obtained. For quantification of mng number in *eyGal4* and *eyGal4>Nerfin-1 RNAi* brains, mng marked by Repo were counted on representative focal planes in 5 brains per genotype, and the p value is calculated using student t-test.

### Data Availability

The raw and processed scRNA-seq data have been deposited to GEO, and the accession number is GSE168553.

## Acknowledgments

We thank the Flow Cytometry Facility and the DNA Services Laboratory, and the High Performance Computing in Biology Group of the Roy J. Carver Biotechnology Center at the University of Illinois at Urbana-Champaign for FACS sorting, and for construction and sequencing of the 10x V3 Single Cell libraries, and initial analysis of the sequencing data, respectively, and for providing the corresponding method part. We thank the fly community, especially Claude Desplan, Steven Russell, Cheng-Yu Lee, Tzumin Lee, Deborah Hursh, J. Peter Gergen, Tiffany Cook, Adrian Moore, Chris Doe; Richard Mann, Andrew Tomlinson, John R. Nambu and Makoto Sato, for generous gifts of antibodies and fly stocks. We thank the Bloomington Drosophila Stock Center, the Vienna Drosophila RNAi Center, the Developmental Studies Hybridoma Bank, and TriP at Harvard Medical School (NIH/NIGMS R01-GM084947) for fly stocks and reagents. We would like to thank the NSF-Simons Center for Quantitative Biology at Northwestern University for supporting this project as a Pilot grant, and also the National Eye Institute for grant support (Grant 1 R01 EY026965-01A1).

## Author contributions

X.L. and H.Z. designed the project and experiments, S.D.Z. analyzed sc-RNA-seq data, H.Z. and X.L. performed experiments and analyzed data, A.R. and Y.Z. participated in some experiments and data discussion. The manuscript is written by H.Z., X.L. and S.D.Z., with all authors commented.

## Materials & Correspondence

Publicly available fly lines (those from BDSC or VDRC) should be requested directly from the corresponding stock centers: https://bdsc.indiana.edu/ or https://stockcenter.vdrc.at/control/main. Fly lines generated in this study can be requested without restriction. Correspondence and requests for materials should be addressed to Xin Li.

---

**Supplementary Figure S1.**
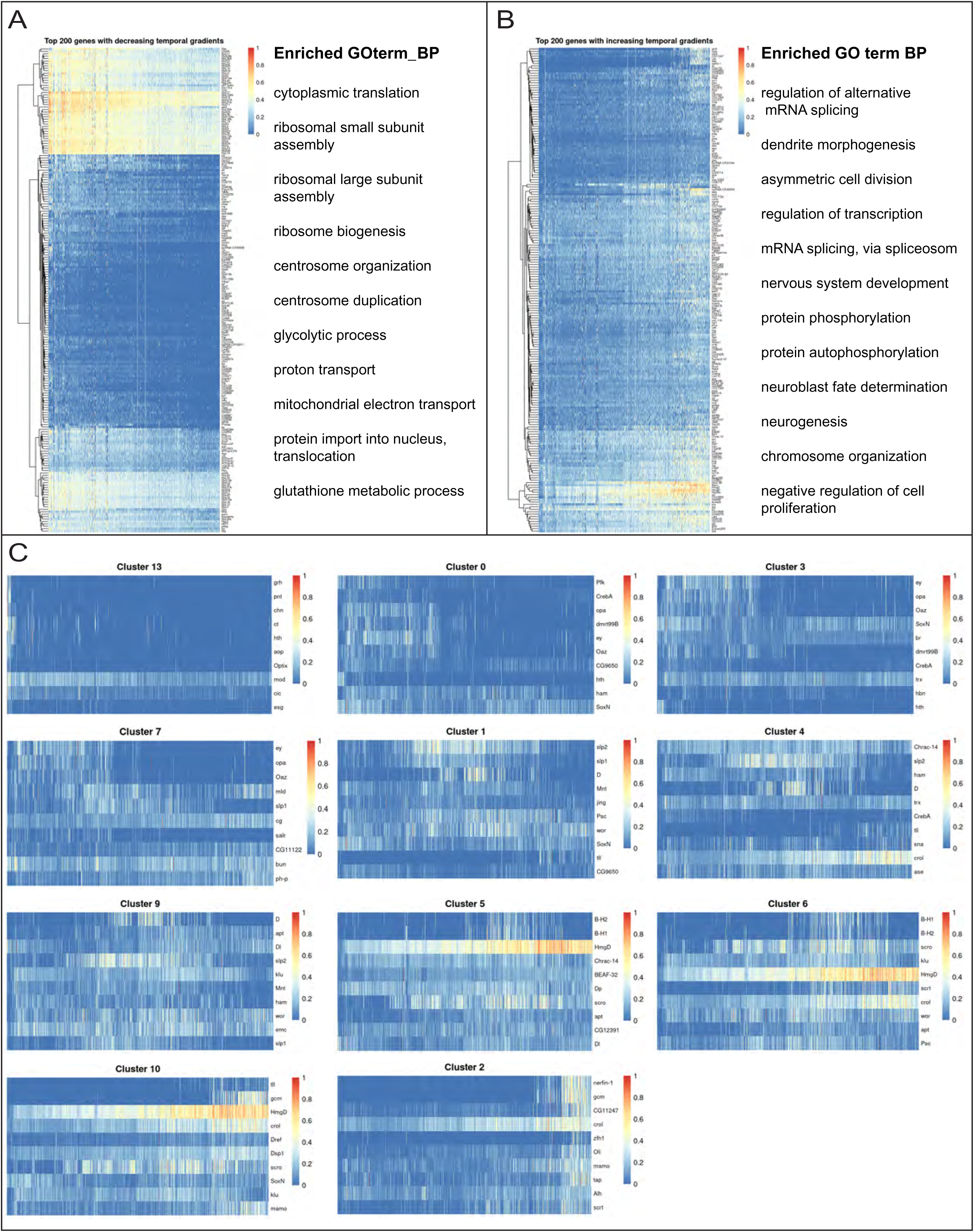
Analysis of genes that show a temporal gradient of expression and TFs differentially expressed in each cluster. In all heatmaps, the expression levels are visualized as a percentage of the maximum observed expression across all cells. (A) Heatmaps showing the top 200 genes with decreasing temporal gradients across the pseudotime. Using the Functional Annotation Tool from the DAVID Bioinformatics Resources 6.8, these genes are enriched in the GO terms related to translation and metabolism. For details refer to Supplementary Table 2. (B) Heatmaps showing the top 200 genes with increasing temporal gradients across the pseudotime. These genes are enriched in the GO terms related to the regulation of gene expression, neural development, and signal transduction among others. For details refer to Supplementary Table 3. (C) Heatmaps showing the top 10 differentially expressed TFs for each NB cluster. The clusters are ordered according to their positions along the pseudotime.

**Supplementary Figure S2.**
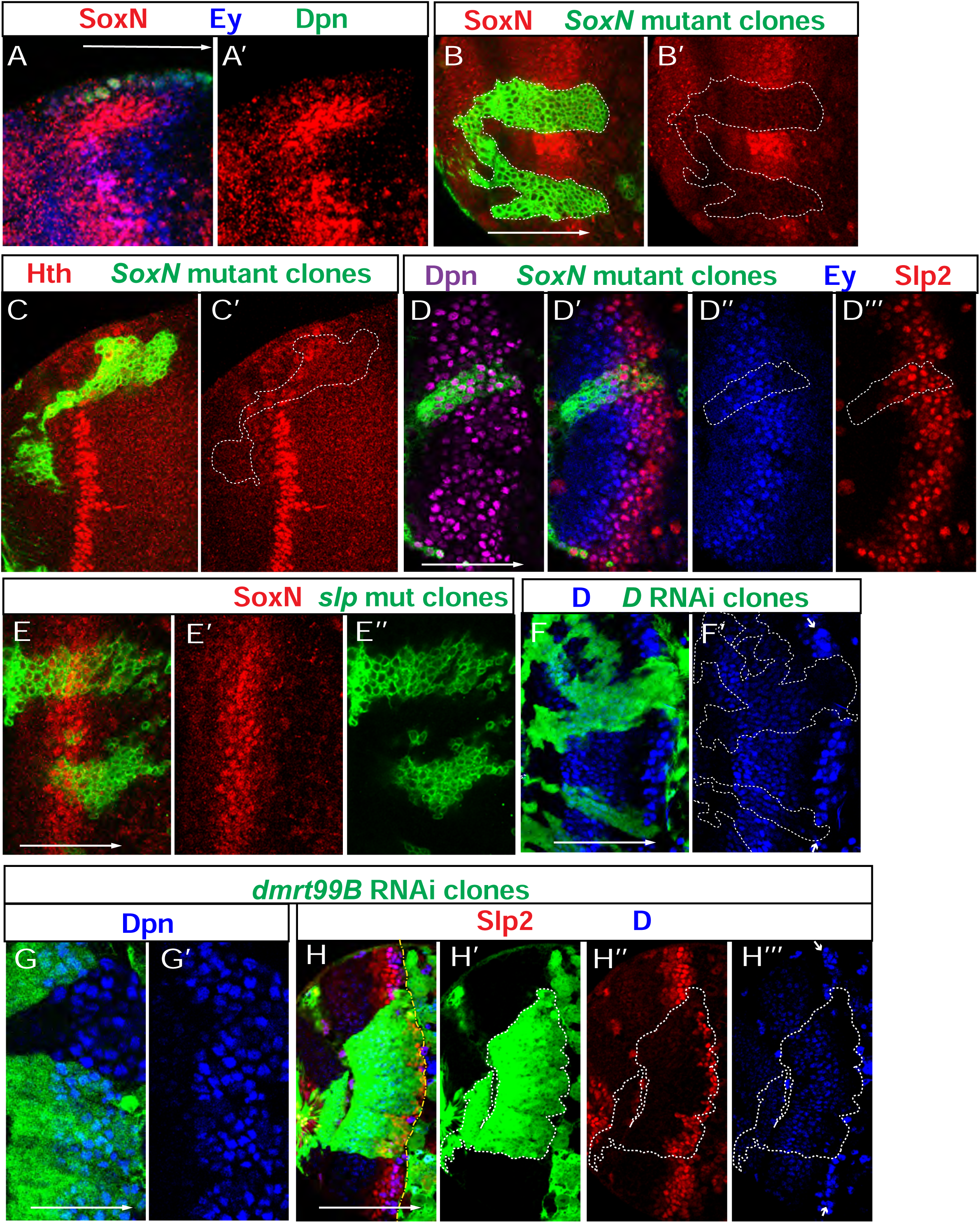
Additional data for SoxN and Dmrt99B. (A-A’’) SoxN (red) is expressed in progeny generated before the Ey (blue) expressing progeny. NBs are marked by Dpn (green). (B,B’) In *SoxN^NC14^* mutant clones marked by GFP (green), SoxN (red) is lost (6 out of 6 clones). (C,C’) In *SoxN^NC14^* mutant clones marked by GFP (green), Hth expression (red) is normal (5 out of 5 clones). (D-D’’’) In *SoxN^NC14^* mutant clones marked by GFP (green), Ey (blue) and Slp2 (red) are still expressed in NBs marked by Dpn (magenta) (9 out of 9 clones). (E-E’’) In *slp* mutant clones marked by GFP (green), SoxN expression (red) is not affected (7 out of 7 clones). (F,F’) The antibody for D (blue) has unspecific cross-reactivity to other proteins in early NBs (the weak staining that is not lost in *D*-RNAi clones), while the second stripe of strong staining (white arrows) is specific for D, as the staining is lost in D RNAi clones (green). (G,G’) In *Dmrt99B-RNAi* clones marked by GFP (green), NB formation is not affected as indicated by Dpn staining (blue) (11 out of 11 clones). (H-H’’’) In *Dmrt99B-RNAi* clones marked by GFP (green), Slp expression (red) is delayed, and D expression (strong blue staining indicated by white arrows) is lost (in 8 out of 8 clones). Note the weak blue staining in young NBs is cross-reactivity against other proteins. The white dashed lines in H’-H’’’ indicate clone margins, while the yellow dashed line with alternating short and long dashes in H indicates the boundary between the medulla and the central brain.

**Supplementary Figure S3.**
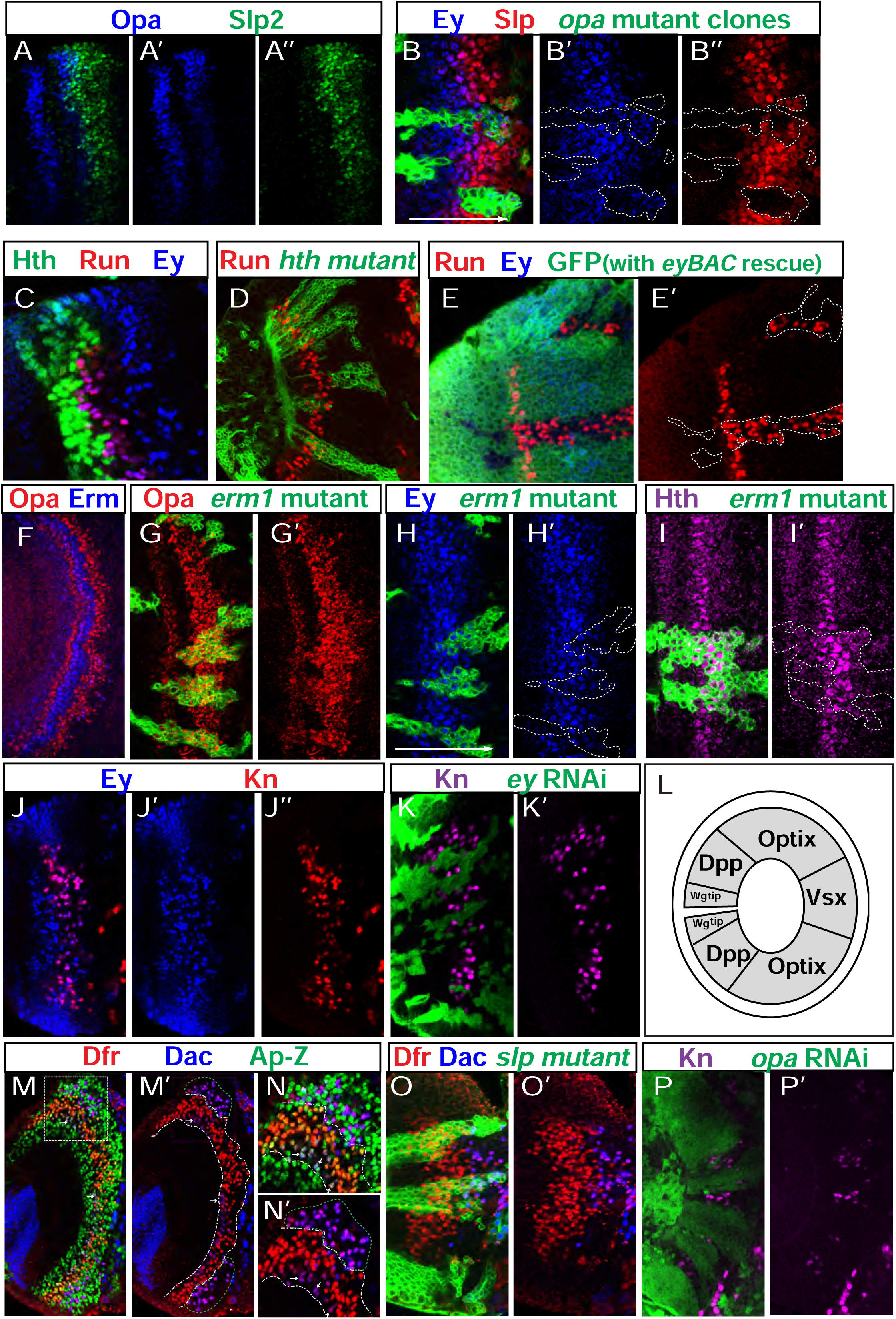
Additional data for Opa and Erm. (A-A’’) Opa (blue) is expressed in two layers of progeny, but is lost as neurons mature. Slp2 expression is in green. (B-B’’) In *opa^7^* mutant clones (marked by GFP in green), the expression of Ey (blue) and Slp (red) is delayed (in 14 out of 14 clones). (C) Runt (red) expressing neurons are generated between the Hth (green) neurons and the Ey (blue) stage neurons. Note: Runt neurons later turn on Ey expression but they are not generated in the Ey stage. (D) In *hth^P2^* mutant clones marked by GFP (green), Runt (red) expressing neurons are still generated. (E,E’) In *ey* mutant clones marked by lack of GFP (green), Ey (blue) is lost, Runt (red) is expanded into later-born progeny. These negatively marked clones are generated in *ey* mutant background, and GFP positive cells contain the *eyBAC* rescue construct, while GFP negative cells are *ey* mutant (enclosed by white dashed lines). (F) Erm::V5 (blue) is expressed in a layer of progeny generated between the two layers of Opa (red) expressing progeny. (G,G’) In *erm^1^* mutant clones marked by GFP (green) at a deeper progeny focal plane, the gap of Opa (red) expression in the progeny is also lost (in 13 out of 13 clones). (H,H’) In *erm^1^* mutant clones marked by GFP (green) at the surface NB focal plane, Ey expression (blue) is still present but weaker (in 10 out of 10 clones). (I,I’) In *erm^1^* mutant clones marked by GFP (green), Hth expression (magenta) is slightly expanded (in 5 out of 5 clones). (J-J’’) Kn (red) is expressed in the Ey-expressing (blue) neurons generated by the Ey stage NBs. Note: A layer of earlier-born Ey expressing neurons (which are the Runt neurons that turn on Ey expression later), do not express Kn. (K,K’) In *ey-RNAi* clones marked by GFP (green), Kn (magenta) neurons are lost (in 10 out of 10 clones). (L) The schematic model of the compartmentalization of the medulla. (M-N’)The expression of Dfr (red), Dac (blue), and *ap-*LacZ (green) in neurons: a layer of earlier-born neurons express both Dfr and *ap-*LacZ (between the two white dashed lines in M’-N’), and several clusters of later-born neurons express both Dfr and Dac but not *ap-*LacZ (enclosed in green dashed circles). White arrows point to cells expressing Dfr, *ap-*LacZ and a weak level of Dac within the first population. (N,N’) A zoomed-in image of the outlined square in M. (O,O’) In *slp* mutant clones marked by GFP (green), Dfr (red) and Dac (blue) expressing neurons are still generated (10 out of 12 clones). (P,P’) In *opa-RNAi* clones marked by GFP (green), Kn (magenta) expression is lost (in 19 out of 20 clones).

**Supplementary Figure S4:**
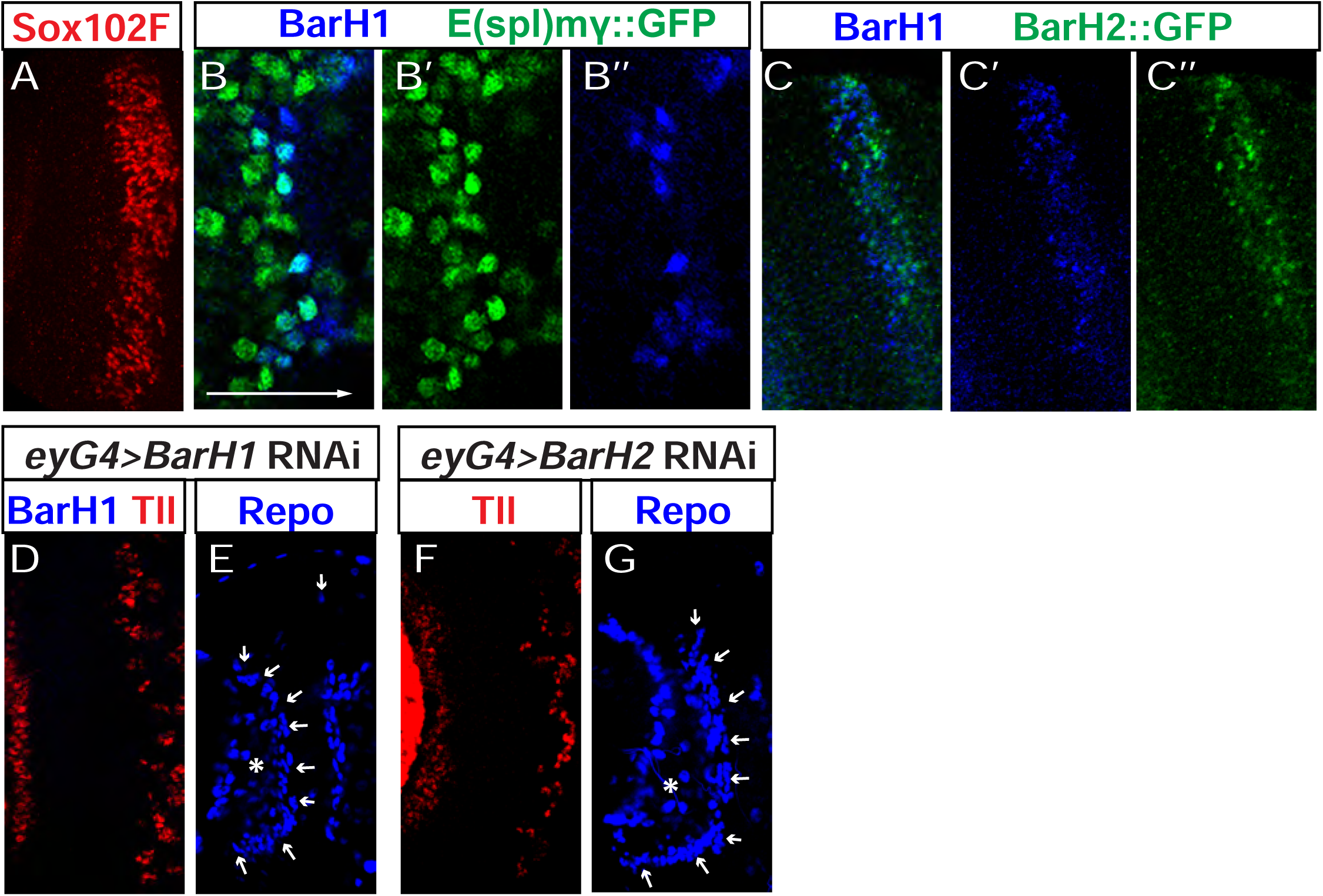
Additional data for Scro and BarH. (A) The expression pattern ofSox102F (red) in neurons. (B-B’’) The expression of BarH1 (blue) in old medulla NBs marked by E(spl)mγ::GFP (green). E(spl)mγ::GFP is expressed in all NBs. (C-C’’) The expression of BarH1 (blue) and BarH2::GFP(green) in neurons at a deep progeny focal plane. (D) At the surface NB focal plane, Tll (red) expression in NBs is not affected when *BarH1* RNAi is driven by *eyGal4* (in 4 out of 4 brains). (E) At a deep progeny focal plane, glia marked by Repo (blue) are not affected (in 4 out of 4 brains) when *BarH1* RNAi is driven by *eyGal4*. The asterisk indicates the center of the neuropil and the arrows are pointing at the glia. (F) At the surface NB focal plane, Tll (red) is not affected in NBs when *BarH2* RNAi is driven by *eyGal4* (in 3 out of 3 brains). (G) At a deep progeny focal plane, glia marked by Repo (blue) are not affected when *BarH2* RNAi is driven by *eyGal4* (in 3 out of 3 brains). The asterisk indicates the center of the neuropil, and the arrows are pointing at the glia.

**Supplementary Figure S5:**
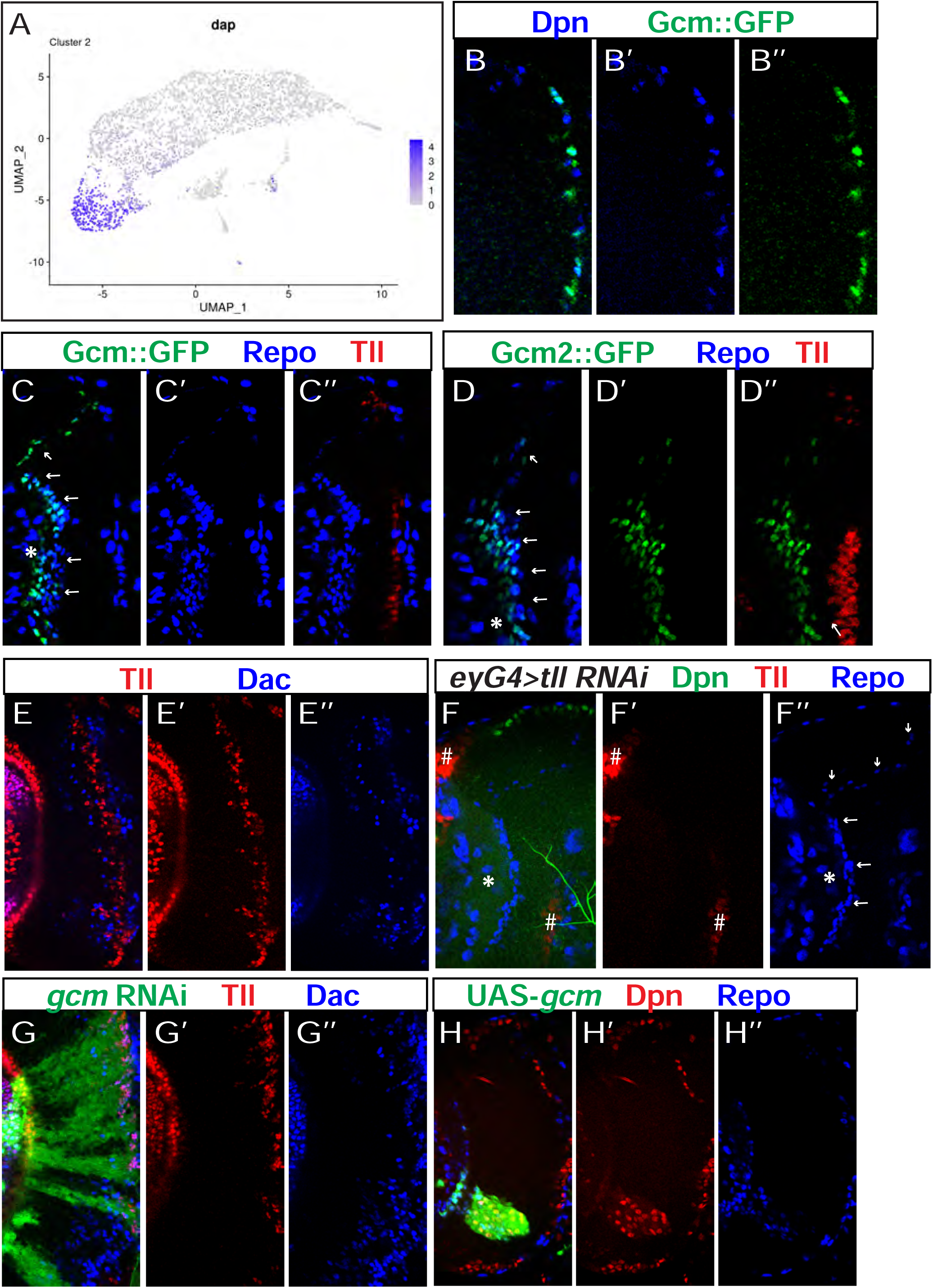
Additional data for Gcm. (A) The mRNA expression pattern of Dap from the scRNA-seq data visualized on UMAP Plot. (B-B’’) The co-expression of Gcm::GFP (green) and Dpn (blue) in the oldest NBs, located between the medulla and the central brain, on a slightly deeper focal plane than younger NBs. (C-C’’) The expression of Tll (red), Gcm::GFP (green) and Repo (Blue) in a cross-sectional view. The asterisk indicates the center of the neuropil, and arrows point to mng. (D-D’) The expression of Tll (red), Gcm2::GFP (green) and Repo (blue) in a cross-sectional view. The asterisk indicates the center of the neuropil, and arrows point to mng. (E-E’’) At a relatively superficial progeny focal plane close to the surface NBs, some Tll (red) and Dac (blue) double positive neurons are present in the wild-type brain. (F-F’’) When *tll* RNAi (*tll*-miRNA) is driven by *eyGal4*, Tll (red) is lost in NBs marked by Dpn (green), while glia marked by Repo (blue) are still aligned continuously around the medulla neuropil (in 3 out of 3 brains). The asterisk indicates the center of the neuropil, and arrows are pointing at the glia. The “#” symbol indicates Tll expression in the NE of OPC and IPC is not affected because eyGal4 is not expressed there. (G-G’’) In *gcm* RNAi clones (marked by GFP in green), Tll (red) and Dac (blue) double positive cells are increased at a deep progeny focal plane (in 16 out of 16 clones). (H-H’’) In *gcm* misexpression clones (green), ectopic NBs marked by Dpn (red) are present, and ectopic glia marked by Repo (blue) are also present (in 2 out of 2 clones).

**Supplementary Figure S6:**
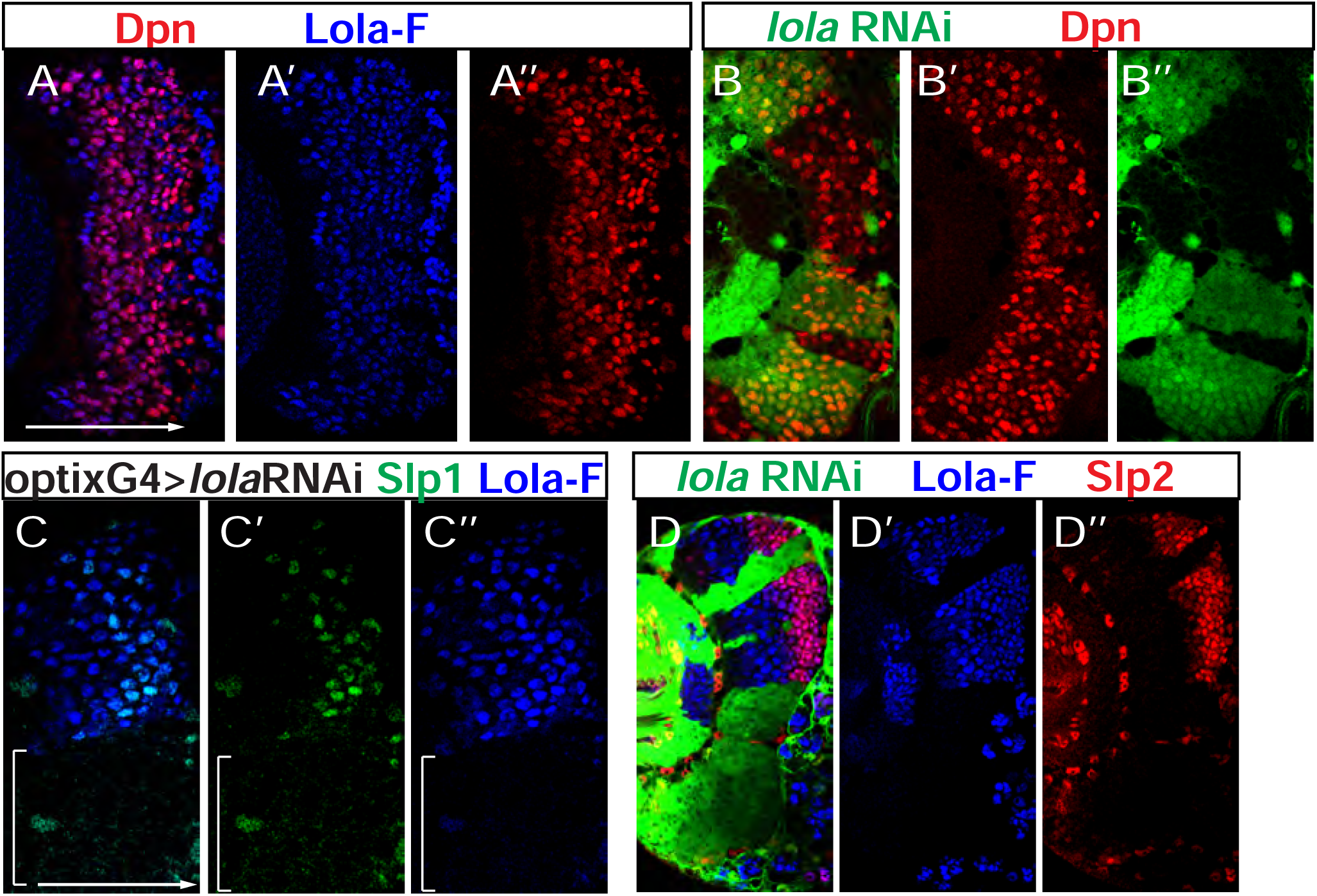
Additional data for Lola and Nerfin-1. (A-A’’) The expression of Dpn (red) and Lola-F (blue) at a surface focal plane. (B-B’’) In *lola* RNAi (BDSC35721) clones marked by GFP in green, the formation of NBs (marked by Dpn in red) from NE cells is not affected. (C-C’’) When *lola* RNAi (BDSC35721) is driven by *optixGal4*, the expression of Lola (blue) is lost and the expression of Slp1 (green) is greatly delayed (in 4 out of 4 brains), in mOPC indicated by a white bracket. (D-D’’) In *lola* RNAi (VDRC 101925) clones marked by GFP (green), the expression of Lola (blue) is lost and the expression of Slp2 (red) is greatly delayed or lost (11 out of 11 clones).

**Supplementary Table 1. Known *Drosophila* cell cycle genes from Tinyatlas at Github (source:** https://github.com/hbc/tinyatlas/blob/master/cell_cycle/Drosophila_melanogaster.csv). This table includes known *Drosophila* cell cycle genes and the corresponding phase, and were used to estimate the cell cycle phase of each cell in our scRNA-seq data.

**Supplementary Table 2. The top 200 genes showing decreasing gradients of expression and the enriched GOTERM analysis.** Genes with temporally patterned expression gradients were identified as those whose expression levels showed significant Spearman correlation with the inferred pseudotime at an FDR cutoff of 0.05. These genes were sorted according to the “corr” column (Spearman correlation), and top 200 genes with the highest negative values were analyzed for enriched Go terms for Biological Processes “GOTERM_BP_DIRECT” using the “Functional Annotation Chart” at the DAVID Bioinformatics Resources 6.8 website (https://david.ncifcrf.gov/home.jsp).

**Supplementary Table 3. The top 200 genes showing increasing gradients of expression and the enriched GOTERM analysis.** Genes with temporally patterned expression gradients were sorted according to the “corr” column (Spearman correlation), and top 200 genes with the highest positive values were analyzed for enriched Go terms for Biological Processes “GOTERM_BP_DIRECT” using the “Functional Annotation Chart” at the DAVID Bioinformatics Resources 6.8 website (https://david.ncifcrf.gov/home.jsp).

**Supplementary Table 4.** List of reagents used in this study with sources and Identifier numbers.

